# fastman: A Fast Algorithm for Visualising GWAS Results Using Manhattan and Q-Q Plots in R

**DOI:** 10.1101/2022.04.19.488738

**Authors:** Soumya Subhra Paria, Sarthok Rasique Rahman, Kaustubh Adhikari

**Affiliations:** School of Mathematics & Statistics, The Open University, Walton Hall, Milton Keynes, MK7 6AA, UK; Department of Medical Statistics, London School of Hygiene and Tropical Medicine, Keppel Street, London, WC1E 7HT; Department of Ecology & Evolutionary Biology, Princeton University, Princeton, New Jersey, 08540, USA; Department of Genetics, Evolution and Environment, and UCL Genetics Institute, University College London, Gower Street, London, WC1E 6BT, UK

**Keywords:** GWAS, Manhattan plots, Q-Q plots, non-model organisms, large data handling

## Abstract

Visualisation of GWAS summary statistics, specifically P-values, as Manhattan plots are widespread in GWAS publications, and many popular software tools are available, such as the `R` package qqman. However, there is a substantial need for further development, such as handling datasets from non-model organisms. We provide a new `R` package, fastman, with major additional capabilities compared to currently available packages. For example, it handles datasets involving genomes from non-model organisms, even at a draft stage consisting of numerous contigs and scaffolds (shorter stretches of assembled DNA sequences) that have not been compiled into chromosomes. Non-numeric chromosome IDs are also supported. Additionally, our package has the capability of plotting other genetic scores, such as other GWAS statistics (e.g. regression betas or odds ratios), fixation index(*F*_*ST*_), D-statistics, and various selection statistics, such as PBS. Importantly, negative or two-tailed values are supported in this package. In our package, we implement a heuristic algorithm that drastically reduces plotting time for huge datasets without losing visual precision, allowing for many different data types and missing data. We also provide substantial additional flexibility in highlighting and annotation. The package can produce plots directly from association outputs by `PLINK`. Alternatively, it can produce plots from any `R` data frame with custom columns and can handle large datasets to generate plots rapidly. It is available for public use at https://github.com/adhikari-statgen-lab/fastman.

## 1. Introduction

In recent years, human genetics has grown immensely in data production and analysis (Uffelmann *et al*. 2021; Visscher *et al*. 2017). A typical chip used to genotype one participant contains half a million to one million genetic variants. Subsequently, imputation – a statistical method to probabilistically infer genotype data of additional variants based on chip genotype information – increases the number of usable variants to 10 million (Visscher *et al*. 2017). Large-scale datasets are emerging from whole genome sequencing as well, such as All of Us Research Program (All of Us Research Program Investigators 2019) data (250k human whole genome), 1000 genomes project (The 1000 Genomes Project Consortium 2015), Genome Sequence Archive for Human (Zhang et al. 2021), and UK Biobank (Sudlow et al. 2015) (500k whole genome human data). Apart from human genetics, cheaper sequencing is also having a huge impact on research concerning traditionally non-model organisms, and large-scale sequencing datasets from non-human organisms are flourishing. Soon, ambitious projects such as the Earth BioGenome Project (Lewin et al. 2018) and its affiliated project networks will produce many of these datasets. Proportional to the rise in the use of big datasets, there has been a rise in the development of fast algorithms for their handling, processing, and visualisation for big datasets and large output files produced by the analyses of them (Chang *et al*. 2015; Kim et al. 2019; Ziegler, Hartsock, and Baxter 2015; Wright and Ziegler 2017). An example is the widespread use of software to visualise genome-wide association study (GWAS) summary statistics (Turner 2018; Hwang 2016).

The objective of GWAS is to understand the association of genotypes with phenotypes. This is performed by testing for differences in the allele frequency of genetic markers between individuals with similar ancestries but different phenotypes. In GWAS, hundreds of thousands of genetic markers across many genomes are tested to find which variants have a statistically significant association with a particular trait or disease of interest. The most popularly studied genetic variants in GWAS are single-nucleotide polymorphisms (SNPs), germline substitutions of single nucleotides at specific positions in the genome. The testing is done by performing multiple linear regressions on SNPs and basic covariates like age, sex, etc. A standard set of GWAS results consists of a list of SNPs, their associated chromosomal position, and a P-value representing the statistical significance of the association of interest.

The Manhattan plot (Figure 1) is one of the most popular ways of visualising GWAS results. This is a plot of the negative logarithms of P-values on the y-axis versus the chromosomal position of the SNP on the x-axis. Hence, each dot in a Manhattan plot represents an SNP. The stronger the association, the smaller the P-value, and the higher the value of the negative logarithm. Hence, SNPs with the most significant associations are positioned at the top of the graph, giving the plot the appearance of a Manhattan skyline, a group of skyscrapers rising above the standard buildings.

**Figure 1:**
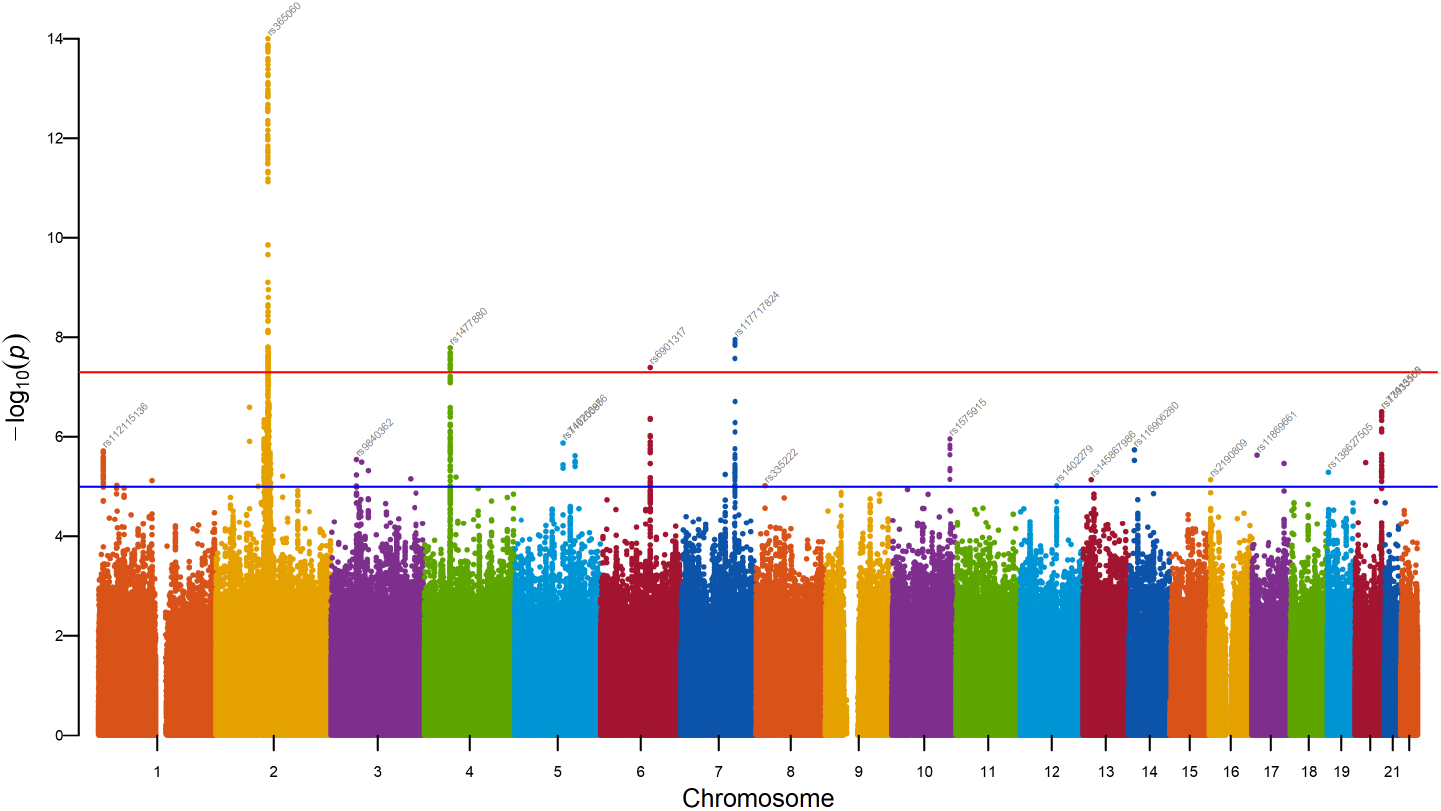
A Manhattan plot.

Another popular plot for viewing GWAS results is the quantile-quantile or Q-Q plot. In the Q-Q plot (as shown in Figure 2), the observed P-values for each SNP are sorted from largest to smallest and plotted against the expected uniform distribution of P-values under the null hypothesis (zero association). If the observed values correspond to the expected values, all points are on or near the diagonal line. Deviation from this line indicates possible inflation (or deflation) of test statistics. This may happen because of population stratification, genotyping errors, or systemic bias in the analysis. One way to quantify inflation is through the **genomic inflation factor** (*λ*), which is defined as:

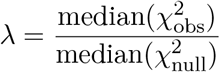

- 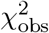 represents the observed test statistics from the GWAS,
- 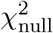 represents the expected test statistics under the null hypothesis (which follows a *χ*^2^ distribution with 1 degree of freedom),
- The median of the 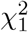 distribution under the null is approximately 0.456.

**Figure 2:**
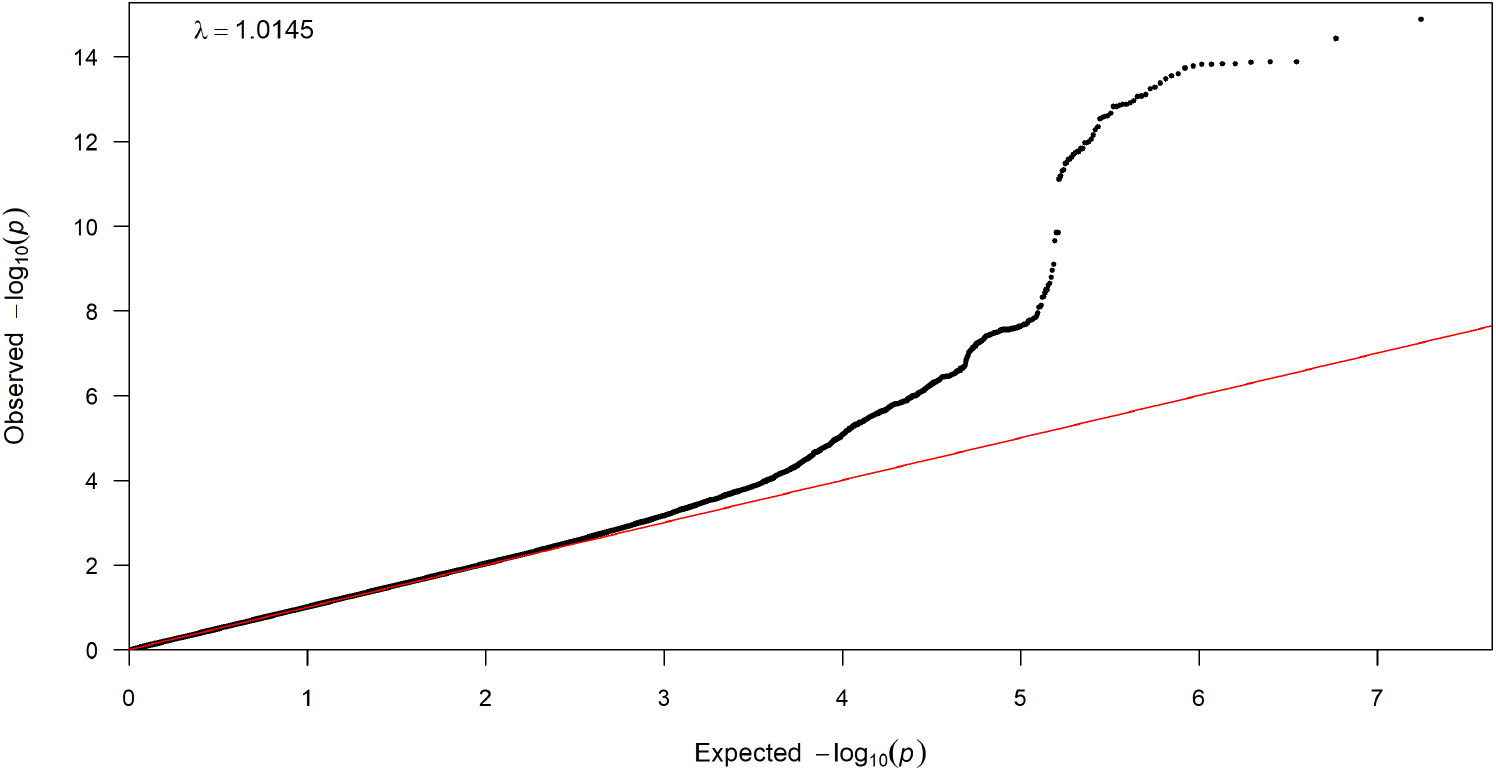
A QQ plot where most of the SNPs follow the expected null distribution but some SNPs on the upper right corner are substantially significant and deviate from the diagonal line.

A *λ* value substantially greater than 1 suggests that test statistics may be inflated, potentially due to uncorrected confounding factors. Conversely, a *λ* value below 1 indicates deflation, which could occur due to overcorrection or conservative testing procedures.

In an ideal scenario, *λ* should be close to 1, which indicates that test statistics follow the expected null distribution. However, in large GWAS with millions of samples and SNPs, slight inflation is common due to polygenic effects rather than systematic bias. Therefore, interpreting Q-Q plots alongside *λ* provides a clearer picture of whether observed deviations are biologically meaningful or results of bias in the study design.

Several software and `R` packages are available for visualising GWAS results using Q-Q and Manhattan plots. Table 1 provides a chronological list of these software and R packages.

**Table 1:**
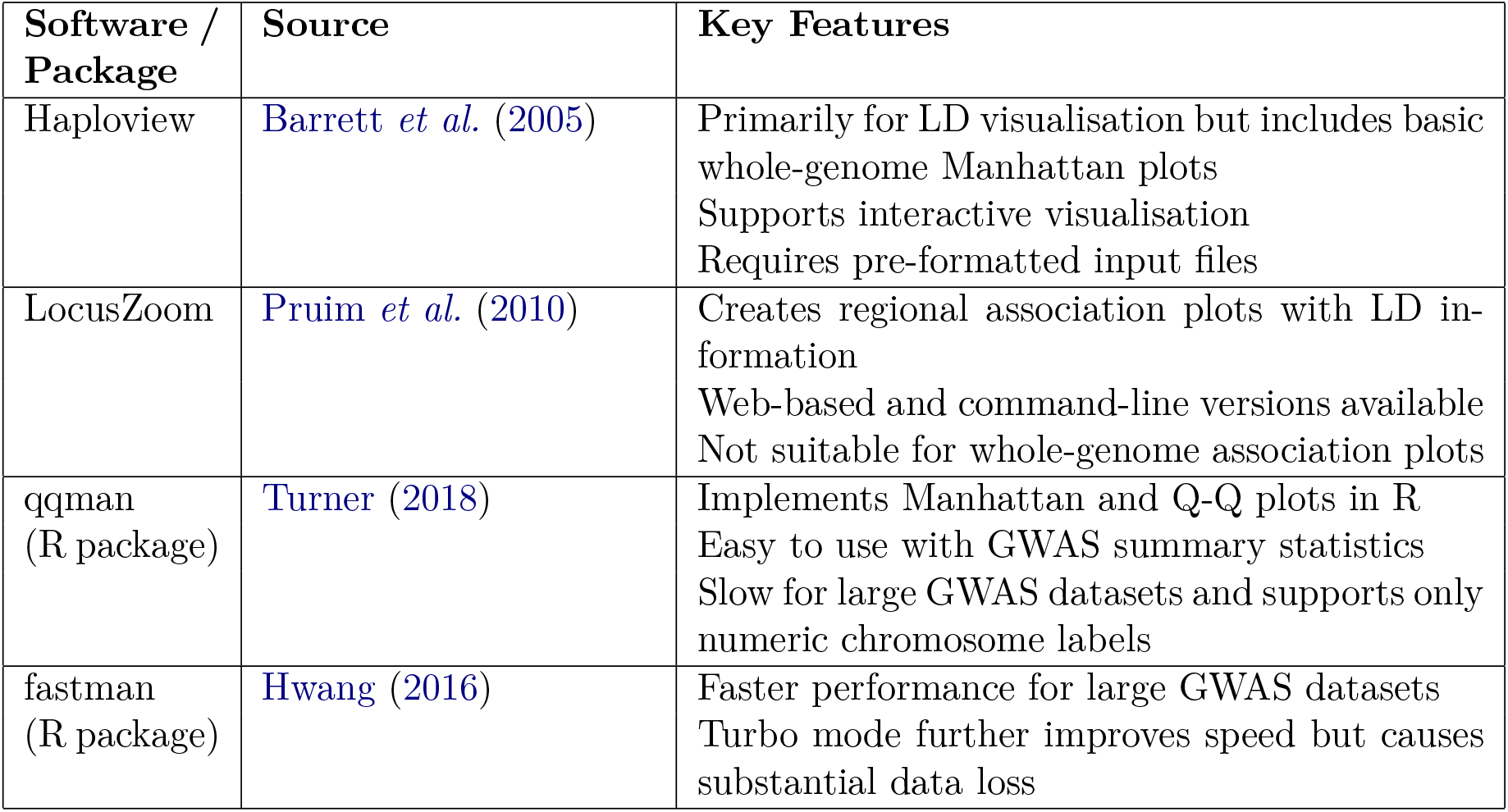
A Chronological List of GWAS Visualisation Software and R Packages.

| Software / Package | Source | Key Features |
| --- | --- | --- |
| Haploview | Barrett et al. (2005) | Primarily for LD visualisation but includes basic whole-genome Manhattan plots<br>Supports interactive visualisation<br>Requires pre-formatted input files |
| LocusZoom | Pruim et al. (2010) | Creates regional association plots with LD information<br>Web-based and command-line versions available<br>Not suitable for whole-genome association plots |
| qqman<br>(R package) | Turner (2018) | Implements Manhattan and Q-Q plots in R<br>Easy to use with GWAS summary statistics<br>Slow for large GWAS datasets and supports only numeric chromosome labels |
| fastman<br>(R package) | Hwang (2016) | Faster performance for large GWAS datasets<br>Turbo mode further improves speed but causes substantial data loss |

Manhattan plots can be created using standalone desktop software `Haploview` (Barrett *et al*. 2005), or for focused regions using the web-based application `LocusZoom` (Pruim et al. 2010). The most popular `R` package for generating the Q-Q and Manhattan plots directly from `PLINK` (Purcell et al. 2007) result files is **qqman** (Turner 2018). **qqman** is an `R` package that contains two functions, `manhattan()` and `qq()`, which can be used to generate the respective plots. Both functions take a data frame with columns containing the chromosome number, chromosomal position, and P-value as input. The typical input should be a `PLINK` .assoc output, but the package allows users to use a custom data frame while specifying the required column names. The default output is a black and white plot with horizontal lines drawn at −log_10_(1× 10^−5^) for “suggestive” associations and −log_10_(5 ×10^−8^) for the “genome-wide significant” threshold. The user has the option to change the colour scheme and to change the threshold levels. The function also gives the option to highlight SNPs of choice if the user provides a column of SNPs. The package also offers a wide range of annotation options for the user, and it being available as an `R` package, the user can use `R` to control every granular aspect to customise the output to their choice.

Despite the plethora of options provided by qqman, it lacks versatility in input handling. qqman only accepts P-values as inputs, and it is incompatible with other genome-wide population genetic parameters like *F*_*ST*_, *π*, and D-statistics (discussed in detail at appendix A). These scores might represent various genetic parameters like degree of association or measure of genetic differences and are very important for several genetic studies. Since these are scores, the ranges and distribution of their values are quite different from that of P-values. As qqman does not support non-numeric entries for chromosome number, results from genomes of non-model organisms cannot be plotted using this package. The genetic sequencing of non-model organisms is becoming increasingly common as advances in sequencing technology enable low-cost, high-quality sequencing. Unlike model organisms such as humans or mice, for which well-annotated reference genomes are available, non-model organisms often lack a standardised reference genome. Instead, their genomes are typically assembled from sequencing small fragments, known as contigs, which are then pieced together to infer genomic structure. Due to the incomplete or draft nature of these assemblies, the data cannot always be neatly organised into chromosomes, as is possible for model organisms. Instead, association results are mapped to specific variants and contigs, which frequently have names that include alphabets, numerals, and symbols. Many visualisation tools, including qqman, are designed to handle chromosome-based data structures and do not support such non-standard naming conventions, and hence are unsuitable for studies involving non-model organisms. Also, qqman cannot handle missing values in the input data frame.

Another drawback of qqman is the time for plot generation. On a typical imputed `PLINK` .assoc file of 8.8 million SNPs, qqman takes 834 seconds to generate a Manhattan plot, a considerable duration when the study needs to be performed on multiple traits or cohorts. Therefore, we have developed a package fastman in `R` for fast and efficient visualisation of GWAS results using Q-Q and Manhattan plots. This package creates the plot directly from .assoc outputs provided by `PLINK`, one of the most popular software used for GWAS. In addition to a standard `PLINK` output, the package can produce the plots from any data frame with chromosome, position, and P-value, and is equipped to handle big data with fast plot generation and optimised memory usage.

Coincidentally, there is another existing package called fastman (Hwang 2016) (we will refer to this package as fastman (Daniel) going forward) which offers far fewer user options and input flexibilities compared to us, but takes a substantially lower amount of time to generate Manhattan plots. The package does so by retaining for plotting at most 20000 SNPs with p-values greater than 0.1, thereby causing data loss before plotting. It further offers a turbo (fast) mode which removes all SNPs with P-values greater than 0.1.

Quite often, `R` users on a cluster have permission issues, which cause difficulties in installing packages. Even when permission is not a problem, there can be compatibility issues between available versions of `R` and package versions, or the availability of other tools such as a `C++` compiler or `Java` or `RJava`, which preclude installation of certain packages such as the very commonly used **tidyverse**. Therefore, one of our key design principles was to provide a vanilla `R` version of the functions that users can upload to their cluster as a text file and invoke with `source()`. At the same time, we recognised the convenience of installed packages for those who can do so, and offered a GitHub installable version. We also acknowledged the widespread use of **ggplot** over base `R` plotting and its modification capabilities, as well as the convenience of integrating with other plotting packages that are compatible; hence, we offer a **ggplot** version of the functions, which can be either sourced or installed similarly. Thus, we hope the package provides a broad enough usability for the diverse range of users, and we welcome further feedback from users on improving the provisions.

## 2. Features and Customisations

**fastman** is an R package for fast and efficient visualisation of GWAS results using Q-Q and Manhattan plots directly from `PLINK` output files. The package contains four functions: `fastman()` and `fastqq()`, and their **ggplot** equivalents `fastman_gg()` and `fastqq_gg()`. The `fastman()` function can be used for visualising Manhattan plots and `fastqq()` for Q-Q plots.

### 2.1 Features and Functionalities

The main features of the package are:

#### Speed

It drastically reduces plot generation time compared to **qqman**, the most popular existing package used for this purpose. On a typical imputed `PLINK` .assoc file of 8.8 million SNPs, plotting time is reduced from 834s in **qqman** to 57s for **fastman**. We expect a similar reduction in other datasets as well. In the Results section, we have provided a detailed speed comparison of fastman with other packages.

#### Efficiency

Memory management has been optimised to reduce space usage during plotting. A detailed comparison has been provided in the Results section.

#### Versatility

It can handle various inputs ranging from P-values and logarithms of P-values to *F*_*ST*_ scores. It is compatible with plotting other genome-wide population genetic parameters (e.g., *F*_*ST*_, *π*, D-statistics), regression betas, and odds ratios. It also has provisions to allow both-sided scores, e.g., scores with negative values.

#### Non-model friendliness

Apart from the GWAS results, our package supports plotting results from genomes of non-model organisms (often with hundreds of contigs or many scaffolds) and provides alphabetical and other ordering options.

#### Flexibility

The package can be installed by using `devtools:install_github()` command in `R`. If the users work in an environment where it is difficult or not permissible to install software packages, they can download and source the necessary components from the GitHub repository (Paria and Adhikari 2024). The package also allows users to employ **ggplot2** versions of the functions. Since the package offers the user an option of returning a **ggplot2** object, users can add further customisations with the **ggplot2** syntax. E.g. this enables the incorporation of ggplot2 aesthetics according to user preferences.

#### Annotation and highlight versatility

It provides the user with a wide set of options to customise annotating and highlighting SNPs of interest. Various illustrations have been provided in the Results section, and many more examples are on the GitHub page.

#### Familiarity

We understand that **qqman** is a popular package for visualising GWAS results. Keeping that in mind, we have kept a very similar set of input arguments and code structure compared to **qqman** to maintain a high degree of familiarity for the user.

#### Missing value handling

**fastman** can handle missing values in the input data frame. It checks the data frame, only in the specific columns relevant for plotting, for the presence of any missing data, and removes those rows before plotting. Packages such as qqman fail to plot datasets with missing data unless the data is pre-processed to remove missingness.

#### Gene annotation capability

Besides the standard annotation options, the package also allows annotation with gene names instead of SNP names if the user prefers so. Currently, the package provides in-built support for human builds GRCh36, 37, and 38, but other builds e.g. for non-model organisms can be provided.

### 2.2 Customisation Options

In addition to the speedup procedure, **fastman** offers users a spectrum of options for plot generation. If the users work in an environment where it is difficult or non-permissible to install packages, they can download the contents manually from GitHub and then use the `source()` function in R to call the R code. Apart from this flexibility, **fastman** also provides a versatile range of functionalities for plot customisation. If the users can download and install packages, they can utilise various packages that work along with **fastman**.

Firstly, users can generate a **ggplot** object instead of base R plots. This can be accomplished using `fastman_gg()` function in the package. The function requires ggplot2 to be installed and loaded. **scattermore** is another package that works smoothly along with **fastman**. Both `fastman()` and `fastman_gg()` functions contain a parameter **scattermore**. This option, if chosen, uses the **scattermore** package to further expedite the plot generation process, contingent on parameter settings, with a predefined default for optimal performance.

Furthermore, users can opt for the `CairoPNG()` command instead of the standard `png()` command, integrating the capabilities of the **cairo** package for plot generation. This alternative proves advantageous in producing high-quality image outputs, including superior graphics and anti-aliasing. However, it necessitates the prior installation and loading of the **cairo** package. Importantly, these options are not mutually exclusive and can be seamlessly combined to achieve intricate plot customisation. As **ggplot2** includes **cairo** capability without loading the package, users can set the `type` parameter to “cairo” within `ggsave()` to accomplish this.

Considering optimal usage practices, users are advised to refrain from saving Manhattan and QQ plots in vector graphic formats such as PDF. Such formats tend to result in larger file sizes, as they preserve individual plotting information for millions of data points, leading to prolonged saving and rendering times. Instead, the recommended practice is to save plots as raster graphics in Portable Network Graphics(PNG) format. This format, characterised by compression without losing quality, provides optimised file size and rendering efficiency.

## 3. Methods

The **fastman** package drastically reduces time in plot generation compared to **qqman**. We achieve this through a speedup algorithm that reduces the data size for fast plotting. Our algorithm for speedup is based on the observation that, given standard parameters for plot visualisation, only a finite amount of visual precision is necessary. For example, in a plot with 100 DPI resolution, any perturbations to the points’ coordinates that are less than 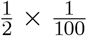 inch in the plot will have no visible effect. Thus, for example, if a range of values 0-10 is being plotted into a 5-inch plot, then a value perturbation of less than 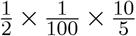 is invisible. This is a conservative calculation because margins take up a part of the plot area, so less than 5 inches will be used to display the values.

We used similar calculations to derive the visual precision required for a high-resolution Manhattan plot with standard dimensions used with lossless export as PNG files. We observed that a Manhattan plot can be perceptually divided into two parts: the top part, where the significant peaks are and where visual accuracy matters most since there are only a few points, and the bottom part, which consists of the non-significant points and is quite crowded therefore visibility of small variations is further limited. Based on the calculations, we concluded that the values plotted along the Y-axis (negative log of P-values) can be rounded to 3 decimal places in the top part and rounded to 2 decimal places in the bottom part, and the resultant plots show no discernible difference when the values are not rounded. Yet, this rounding drastically reduces the number of unique values along the Y-axis. With similar reasoning, we also round the X-axis values, representing chromosomal coordinates, to 2 or 3 decimal places. These rounding steps incur no visible effect on points’ position yet massively reduce the number of unique points, thereby reducing the plotting time, the largest part of the total running time by far. Hence, these rounding steps are the main reason for the speedup in **fastman**, despite the extra computations in rounding and obtaining unique points.

A second heuristic step, elaborated below, finds an appropriate threshold for dividing the Y-axis value into ‘top’ and ‘bottom.’ Due to the flexibility offered by **fastman**, for example, plotting variables such as odds ratios or allele frequencies, in addition to P-values, and also the ability to plot two-sided variables such as regression coefficients, which can be both negative and positive, the heuristic depends on the nature of the variable. It automatically determines whether there is a ‘tail’ on the top or bottom or both or neither side of the plot.

If the data has less than 100k rows, then the full data is rounded to 3 digits, as the rest of the speedup procedure will not be significant in such a small data set. Otherwise, we identify which of the tails of the data are significant, and we round that off to 3 digits. We understand that the data might have a significant right or left tail or both depending on the distribution of the score being plotted. For example, since P-values follow a uniform(0,1) distribution under the null, we expect the negative logarithm of P-values to follow the exponential(1) distribution with a significant right tail. Hence, in the case of P-values, only the right tail is rounded to 3 digits, while the rest of the data is rounded to 2 digits. We use the following statistics to measure the significance of the tails.

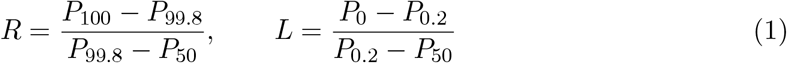

R is used to measure the significance of the right tail, and L is used to measure the significance of the left tail.

*P*_*i*_ here refers to the *i*^*th*^ percentile. The 0.2^*th*^ and 99.8^*th*^ quantiles are heuristics, based on the features of a typical GWAS plot. These statistics are constructed to compare the top and bottom 20,000 of a million to the median. If the data has a significant tail, we expect the corresponding statistics to take a value higher than 0.1. The cut-off value for R and L has been obtained heuristically by looking at various scores and P-value distribution. Figure 3 compares the values of R and L statistics across three distributions: uniform(0,1), normal(0,1), and exponential(1). Bounded parameters such as allele frequency or *F*_*ST*_ can be expected to behave like a uniform distribution, while the negative logarithm of P-values should follow exponential(1). Two-sided data like regression beta are expected to be similar to normal.

**Figure 3:**
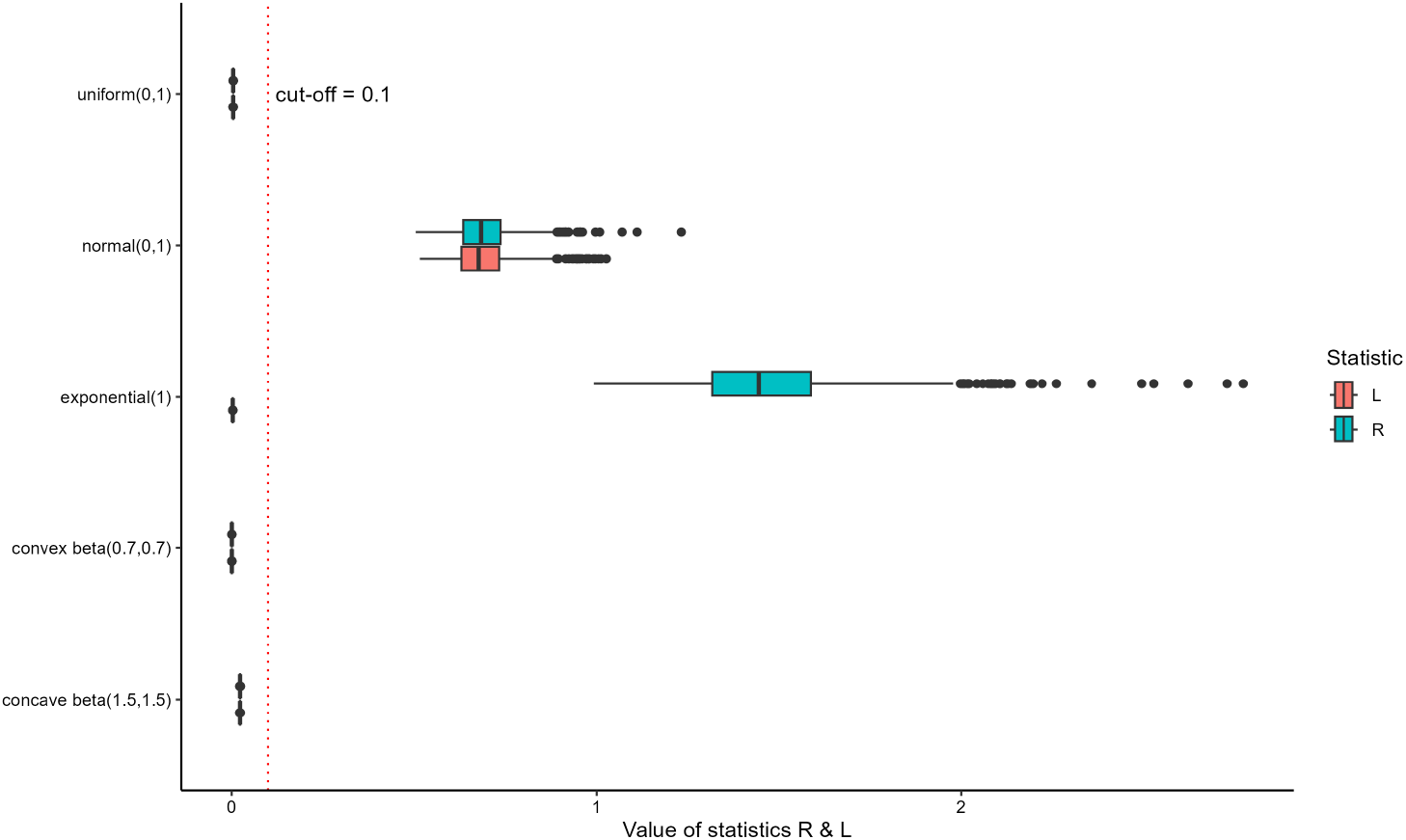
Comparison of statistics R and L across various distributions. The vertical line indicates the 0.1 heuristic cut-off value for R and L to determine the existence of a significant right/left tail in the input data.

Since uniform has no tails, we expect the values of R and L to be less than threshold 0.1. The choice of this threshold has been theoretically validated when P-values follow the null (uniform distribution) (Appendix B), showing that R and L are almost always less than this threshold. Normal has both significant right and left tails; hence, the values of both statistics are above the threshold. For a right-skewed distribution like exponential(1), R is above, and L is below the threshold. The user can cancel the speedup procedure by specifying so in the input parameters. Our speedup heuristic is in contrast to other publicly available Manhattan plot packages that also offer a speedup, such as **fastman (Daniel)**, which reduces the number of points plotted by taking a random subset of the points and is therefore not visually identical to a plot of the original data.

## 4. Results

### 4.1 Illustrations

The **fastman** package includes functions for creating Manhattan and Q-Q plots from GWAS results. Detailed illustrations of its capabilities regarding various input data types and tweaking various visualisation parameters are provided on the package’s GitHub page (Paria and Adhikari 2024).

The following code is for Rstudio users to install the **fastman** package.

~~~
*R> devtools::install_github(‘adhikari-statgen-lab/fastman’,
+        build_vignettes = TRUE)*
~~~

For other users, we would recommend installing the package without building the vignette.

~~~
*R> devtools::install_github(‘adhikari-statgen-lab/fastman’,
+        build_vignettes = FALSE)*
~~~

For the first illustration (Figure 4) of GWAS visualisation, we use data from a GWAS conducted on human participants, probably the most common kind of GWAS. We use a set of published GWAS summary statistics, which were released with the publication of a GWAS (Adhikari et al. 2016) conducted on 6000 participants from various countries of Latin America. The participants were genotyped on a commercial Illumina OmniExpress genotyping chip containing 700K genomic variants. Based on the publicly available reference database called 1000 Genomes, the data was subsequently imputed to increase the number of available genetic variants to 10 million, using the software `Impute2` (Howie et al. 2012). The particular association results used here correspond to the phenotype of facial hair density, which was used here since it contains a clear, prominent association peak on chromosome 2 and other suggestive and genome-wide significant association peaks. So this allows the display of both the suggestive and genome-wide significant association threshold lines and the Y-axis clipping performed by the `maxP` parameter.

**Figure 4:**
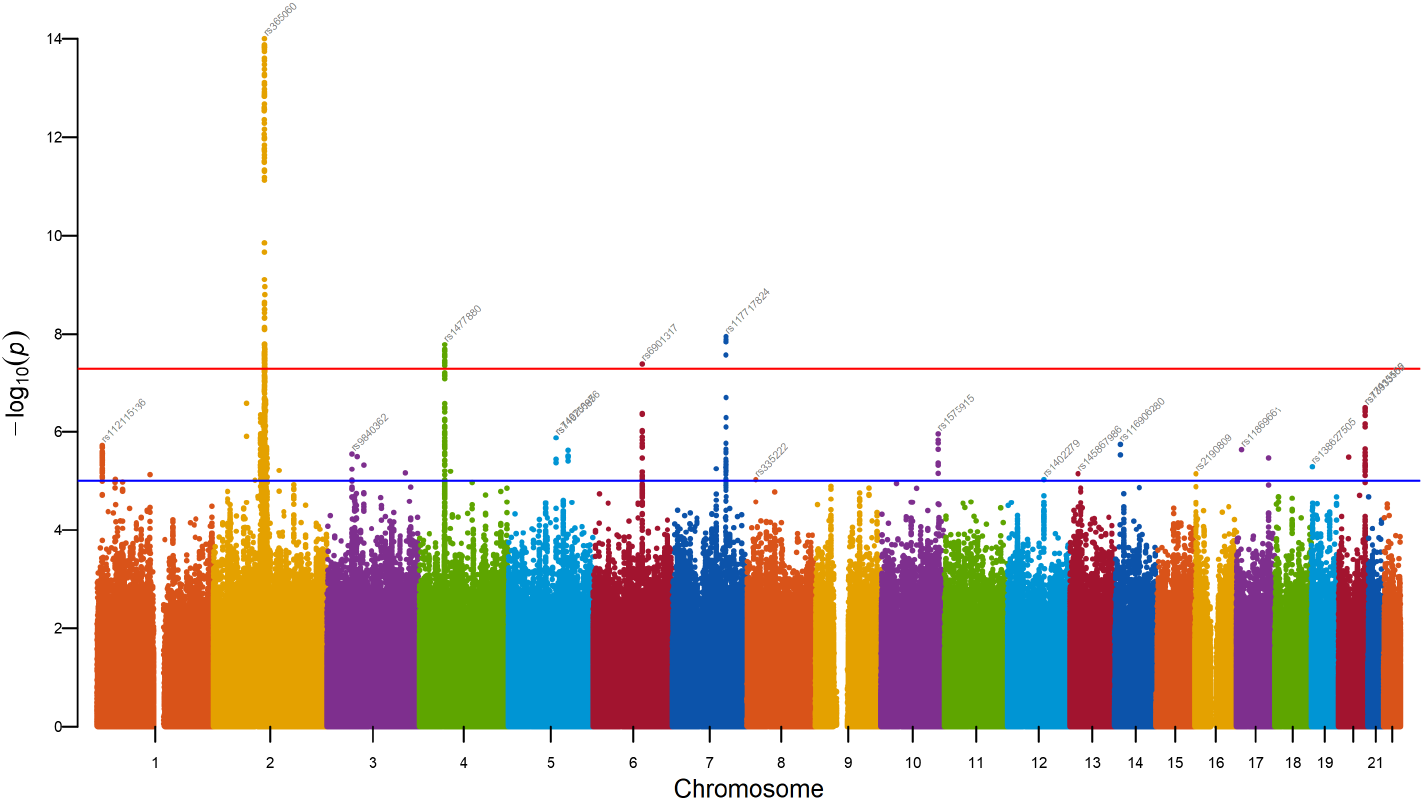
A sample Manhattan P-value plot from `fastman()` using common conventions. Genome-wide significance line (red) and suggestive line (blue) are shown on the plot. Top SNPs from each chromosome have been annotated with a P-value threshold of 1 × 10^−5^.

~~~
*R> data1 <-read*.*table(“beard*.*assoc.linear”, header = TRUE,
+        stringsAsFactors=FALSE, sep = “\t”)
R> str(data1)*
‘data.frame’:        8776423 obs. of 9 variables:
CHR   : num   1 1 1 1 1 1 1 1 1 1 …
SNP   : chr   “rs58108140” “rs180734498” “rs116400033” “rs62637813” …
BP    : num   10583 13302 51479 52058 52185 …
A1    : chr   “A” “T” “A” “C” …
TEST  : chr   “ADD” “ADD” “ADD” “ADD” …
NMISS : num   1551 1911 1552 2348 2421 …
BETA  : num   0.1486 0.07006 0.02572 0.1391 −0.00482 …
STAT  : num   1.793 0.7667 0.3814 2.035 −0.0403 …
P     : num   0.0731 0.4433 0.703 0.042 0.9678 …
~~~

This dataset has results for 8,776,423 SNPs on 22 chromosomes. Let us take a look at the data.

~~~
*R> head(data1)*
 CHR         SNP    BP A1 TEST NMISS      BETA    STAT        P
1  1  rs58108140 10583  A  ADD  1551  0.148600  1.79300 0.07314
2  1 rs180734498 13302  T  ADD  1911  0.070060  0.76670 0.44330
3  1 rs116400033 51479  A  ADD  1552  0.025720  0.38140 0.70300
4  1  rs62637813 52058  C  ADD  2348  0.139100  2.03500 0.04199
5  1 rs201374420 52185  T  ADD  2421 −0.004824 −0.04033 0.96780
6  1 rs150021059 52238  T  ADD  2290 −0.103600 −0.86640 0.38640
~~~

From the above dataset, lets generate a basic Manhattan plot.

~~~
*R>   CairoPNG(“example1.png”, width = 10, height = 6, units = “in”, res = 300)
R>   fastman(data1)
R>   dev.off()*
~~~

For Figure 4, the values of the parameters *R* and *L* are 4.83 and 0.0029 respectively, so only *R* is greater than the threshold 0.1 but not *L*. This confirms that the negative log P-values have a substantial right tail i.e. large positive values, which are thus rounded with higher visual precision. But there is no notable left tail — the small values are part of the bulk of the data that comprise the non-significant P-values.

The package allows the user to choose different annotation colours for different SNPs. The user can only colour the part of the plot above the P-value threshold while the rest of the plot stays grey. The second example (Figure 5) shows a plot using the previous data. We have chosen to annotate the top SNP for every 20-megabase window.

**Figure 5:**
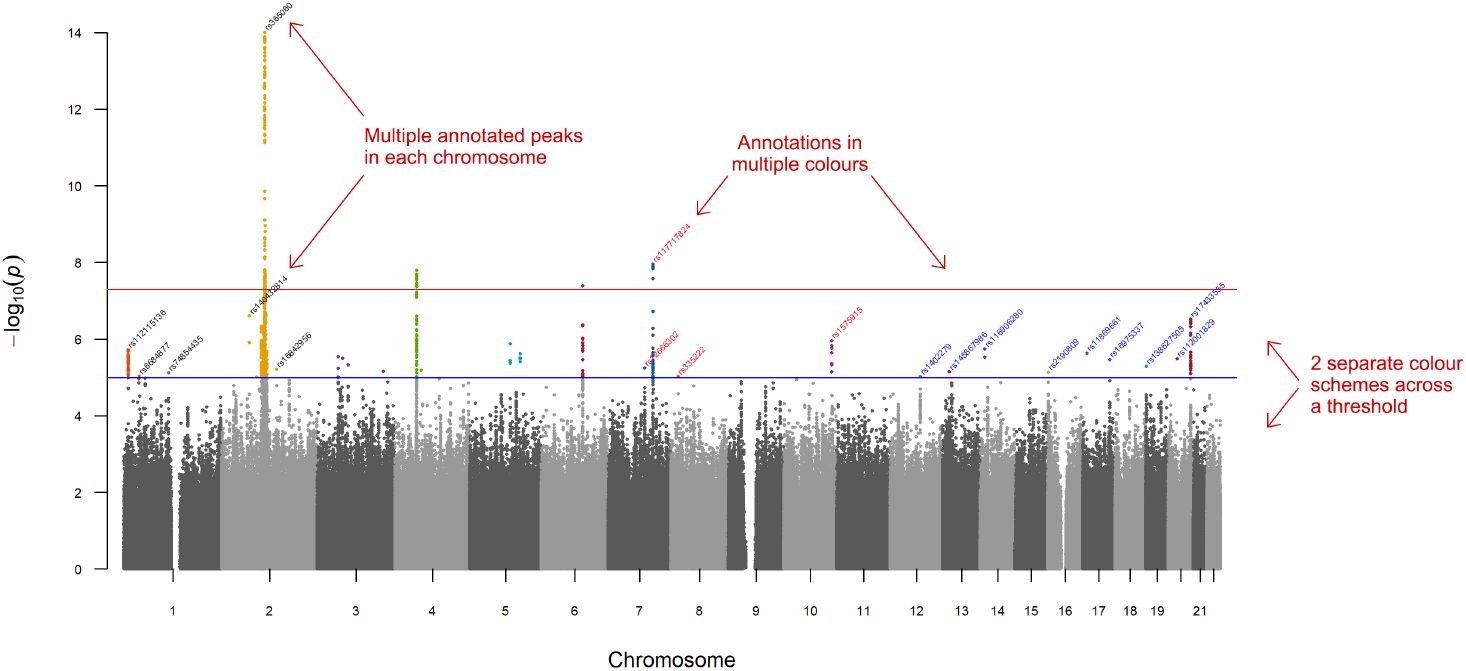
A detailed annotated Manhattan P-value plot from `fastman()`. The descriptions (using arrows) provided in the plot have been manually added for better understanding and have not been generated using the package.

Apart from the logarithm of P-values, scores, including two-sided scores can also be plotted using **fastman**. The next illustration (Figure 6) shows a plot of regression beta coefficients for the GWAS of this quantitative trait. Instead of SNP names, we have chosen to annotate the tops using gene names. For this plot, we are using an annotation threshold of 1.6.

**Figure 6:**
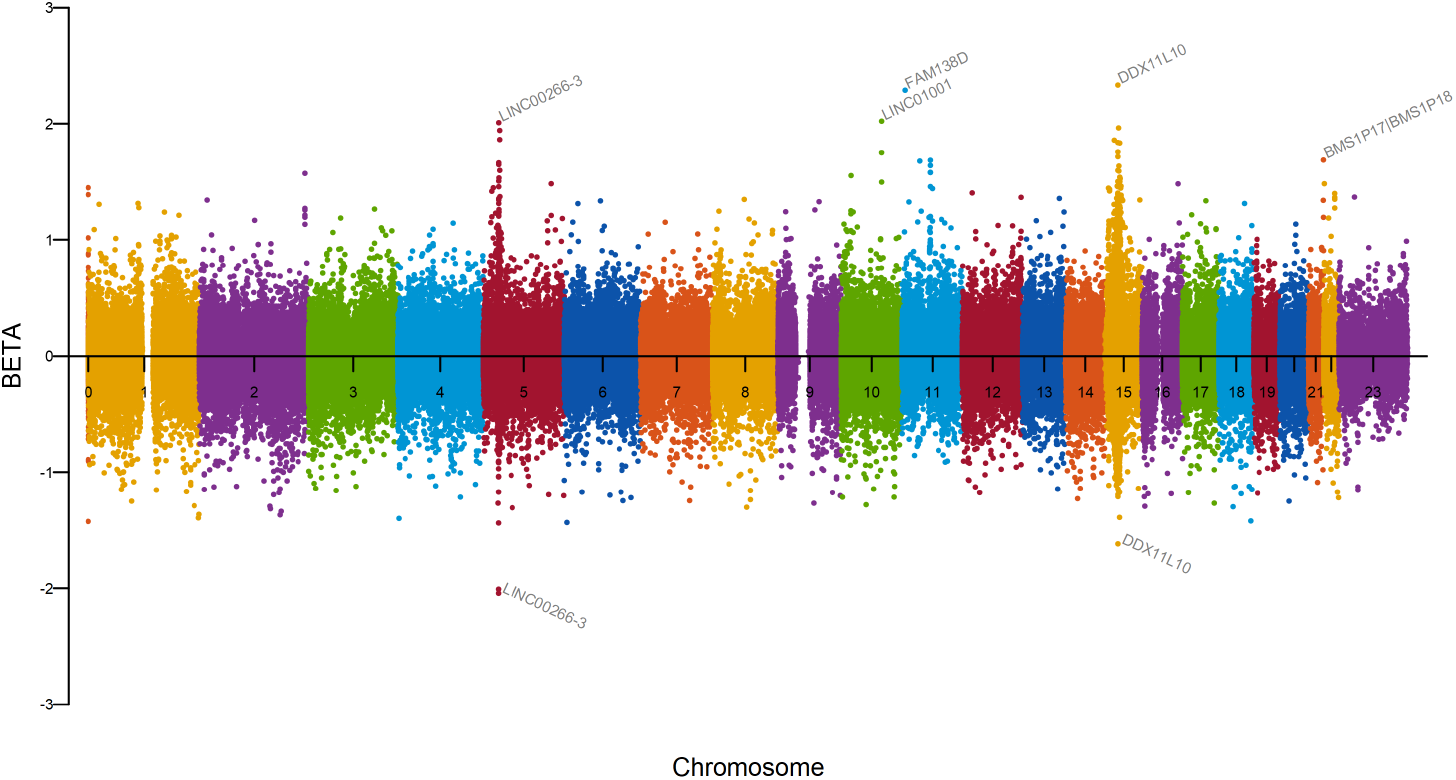
A gene annotated Manhattan beta plot from `fastman()`

For these beta coefficients, consistent with the plot being two-sided, the values of the parameters *R* and *L* are 1.14 and 1.54 respectively, i.e. both values are greater than the threshold 0.1. This confirms that both the right and left tails are notable, and thus rounded with higher visual precision.

~~~
*R>  CairoPNG(“example3.png”, width = 10, height = 6, units = “in”, res = 300)
R>  fastman(data1, p = “BETA”, logp = FALSE, annotatePval = 1.6,
+        geneannotate = TRUE, build = 37, baseline = 0)
R>  dev.off()*
~~~

In the next illustration, we use data from another GWAS (Adhikari et al. 2016) for a binary (dichotomous) trait conducted on the same cohort. A face shape trait of the participants was evaluated on a dichotomous scale to find SNPs with significant associations. Figure 7 shows a plot of odds ratios from the study. The users can add a baseline to their plot at their choice of position, just like the one we have added at the y-axis intercept of 1 in this plot.

**Figure 7:**
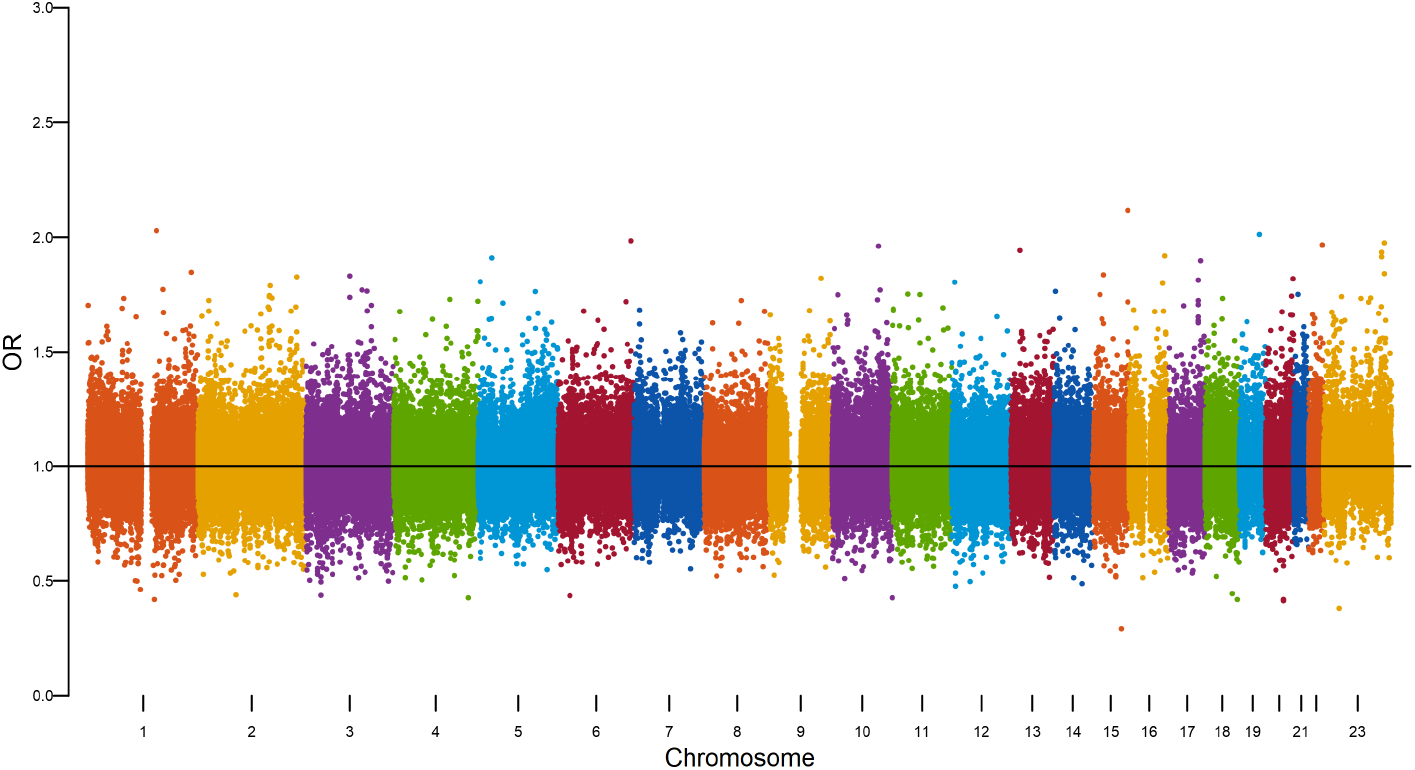
An odds ratio plot with a baseline added at 1 from `fastman()`.

~~~
*R>  CairoPNG(“example4.png”, width = 10, height = 6, units = “in”, res = 300)
R>  fastman(data2, p = “OR”, logp = FALSE, baseline = 1)
R>  dev.off()*
~~~

Most commonly used GWAS visualisation packages like **qqman c**annot handle data from non-model organisms. The genetic sequencing of non-model organisms is becoming popular as new technologies enable low-cost, high-quality sequencing. A challenge of working with non-model organisms is that a standardised reference genome is usually unavailable. To do the genetic sequencing, the genome is divided into small fragments of manageable size, and the fragments are then sequenced. These fragments need to be assembled properly so that the genomic pattern in a region can be established. However, due to the lack of a standardised reference genome, such assembly is usually incomplete or at a draft stage. This means the data cannot be neatly organised into chromosomes, as is possible for humans and other model organisms. Therefore, the small fragments or contigs are directly used instead of chromosomes for analysis and visualisation, so association results refer to specific variants and contigs. Often, a draft assembly will contain thousands of contigs with custom names containing alphabets, numerals, and symbols. Such contig names also cause problems for many software.

For the next two visualisation examples (Figures 8 and 9), we use a SNP data set from the published literature (Rahman *et al*. 2021) that utilised a bumble bee (*Bombus terrestris*) genome assembly (RefSeq GCA_000214255.1; (Sadd et al. 2015); consisting of 10,672 contigs with contig N50 of 76,043 bp) to investigate the genetic basis of a lab-generated (yellow) colour mutant.

**Figure 8:**
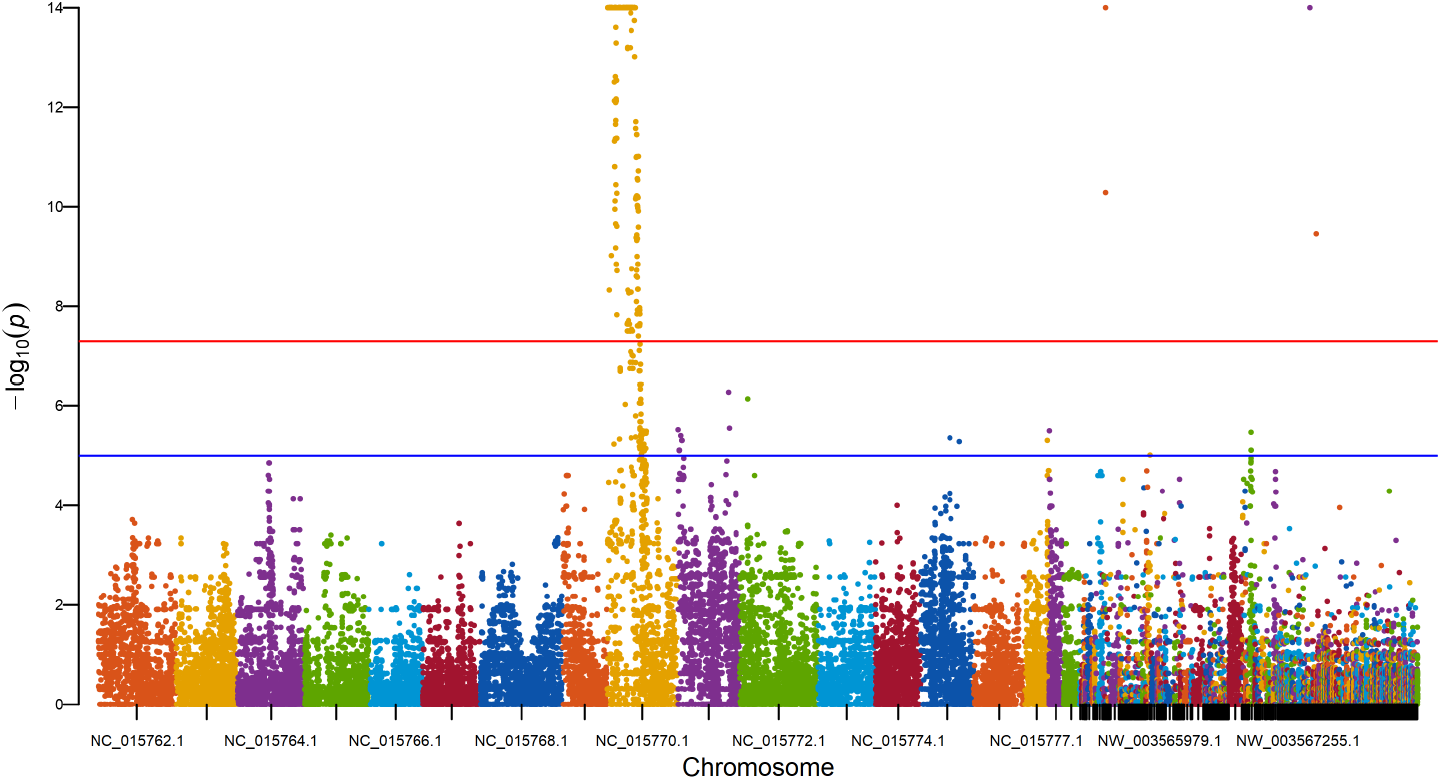
A Manhattan P-value plot from `fastman()` for a non-model organism.

**Figure 9:**
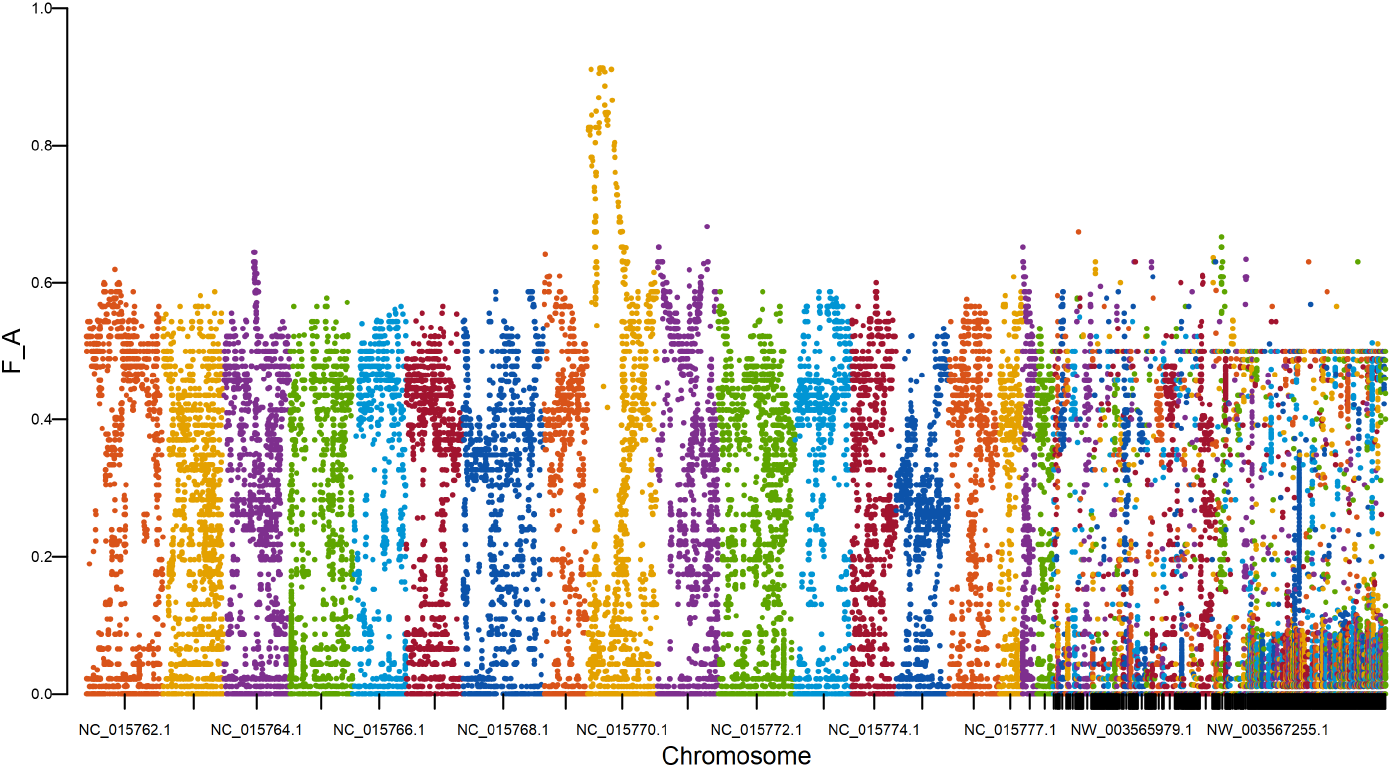
A Manhattan *F*_*ST*_ score plot from `fastman()` for a non-model organism.

~~~
*R> data3 <-read.table(“plink.assoc.fisher”, header = TRUE,
+      stringsAsFactors = FALSE, sep = ‘‘)
R> str(data3)*
‘data.frame’:       57712 obs. of 9 variables:
  CHR:  chr  “NC_015762.1” “NC_015762.1” “NC_015762.1” “NC_015762.1” …
  SNP:  chr  “1139:88:-” “1139:45:-” “1139:28:-” “2518:86:+” …
  BP :  int  32821 32864 32881 58664 68491 83326 83332 92777 92937 92945 …
  A1 :  chr  “T” “C” “G” “T” …
  F_A:  num  0 0.5217 0 0.0222 0.5217 …
  F_U:  num  0.0454 0.375 0.0454 0.0465 0.3636 …
  A2 :  chr  “C” “T” “A” “C” …
  P  :  num  0.0551 0.0528 0.0551 0.436 0.0366 …
  OR :  num  0 1.818 0 0.466 1.909 …
~~~

This dataset has results for 57,712 SNPs on 760 contigs. Let us look at the data.

~~~
*R> head(data3)*
          CHR       SNP    BP A1     F_A     F_U A2       P     OR
1 NC_015762.1 1139:88:- 32821  T 0.00000 0.04545  C 0.05513 0.0000
2 NC_015762.1 1139:45:- 32864  C 0.52170 0.37500  T 0.05276 1.8180
3 NC_015762.1 1139:28:- 32881  G 0.00000 0.04545  A 0.05513 0.0000
4 NC_015762.1 2518:86:+ 58664  T 0.02222 0.04651  C 0.43600 0.4659
5 NC_015762.1 3702:96:- 68491  C 0.52170 0.36360  T 0.03664 1.9090
6 NC_015762.1 4414:33:- 83326  A 0.50000 0.31820  G 0.01556 2.1430
~~~

First, we generate a basic Manhattan plot (Figure 8) for P-values.

~~~
*R>  CairoPNG(“example5.png”, width=10, height=6, units=“in”, res=300)
R>  fastman(data3)
R>  dev.off()*
~~~

The extent of genetic differentiation between two colour morphs (wildtype black vs. mutant yellow) was assessed using the *F*_*ST*_ statistic, a score measuring the amount of differentiation. In Figure 9, we plot the values of this *F*_*ST*_ statistic across the genome to further demonstrate the **fastman** package’s capability of plotting various score statistics, moving beyond just P-values. In the previous examples, the regression beta and odds ratios were unbounded scores, whereas scores such as *F*_*ST*_ or AF (allele frequency) are bounded scores, between 0 and 1.

~~~
*R>  CairoPNG(“example6.png”, width=10, height=6, units=“in”, res=300)
R>  fastman(data3, p = “F_A”, logp = FALSE)
R>  dev.off()*
~~~

The `fastqq()` function accepts a vector of P-values as its input and provides a Q-Q plot with a genomic inflation factor. For this illustration (Figure 10), we use P-values of the data set from our first example. The user can turn off the display of the inflation factor on the plot; it is also returned as the function value.

**Figure 10:**
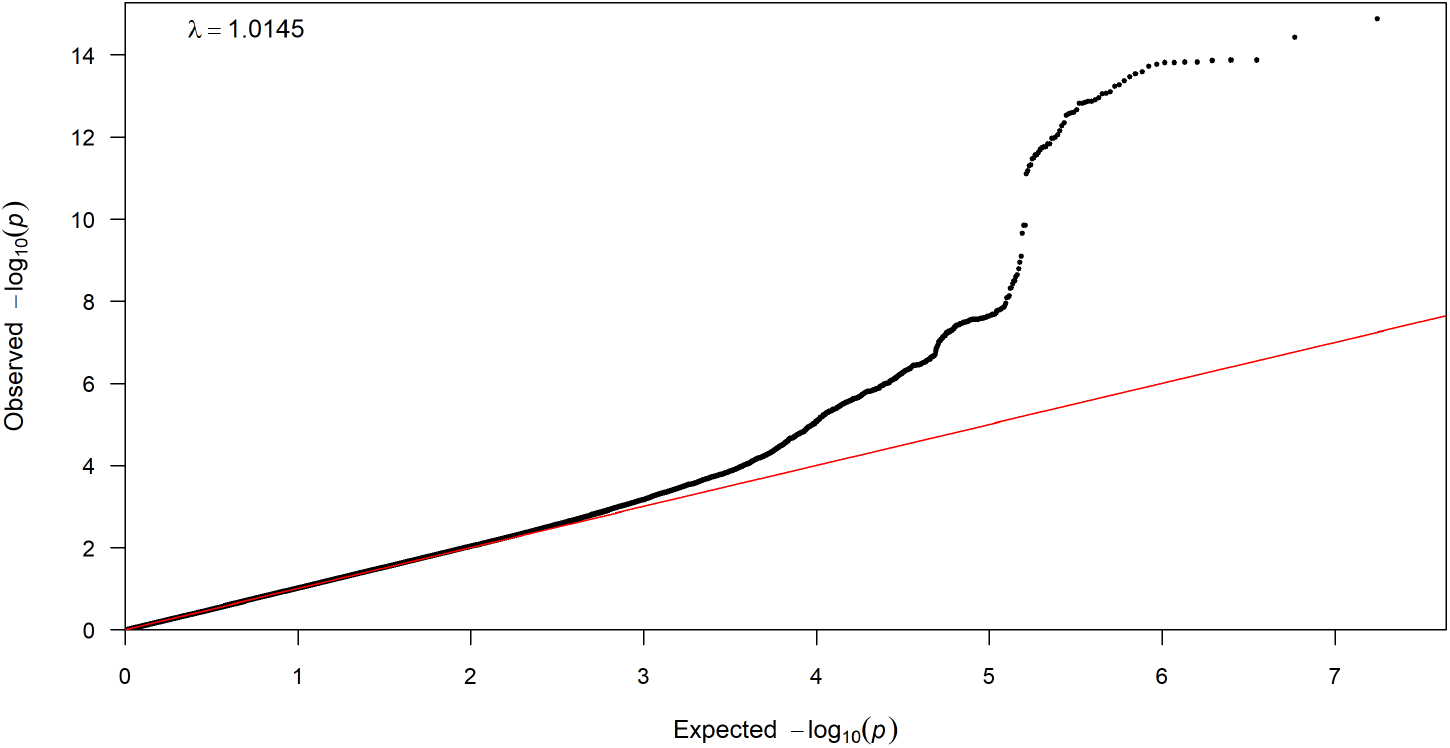
A sample Q-Q plot from `fastqq()`. The red diagonal line represents the reference line (the ideal scenario where the observed P-values exactly match the expected uniform distribution of P-values under the null hypothesis). The genomic inflation factor (denoted by *λ*) has been provided at the top left corner of the plot.

~~~
*R>  CairoPNG(“example7.png”, width=10, height=6, units=“in”, res=300)
R>  fastqq(data1$P)
R>  dev.off()*
~~~

### 4.2. Performance Comparison: fastman

As described in the previous section, **fastman** offers several options for plot generation in addition to the vanilla base R plot generated using the `png()` command. These options can be used in combination with each other. We tried out all possible combinations and compared the time taken for plot generation among all of them. We used the same GWAS summary statistics output file in section 4.1. The data set has results for 8,776,402 SNPs on 22 chromosomes. We ran a hundred iterations on this data set. The box plots of the results are provided in Figure 11.

**Figure 11:**
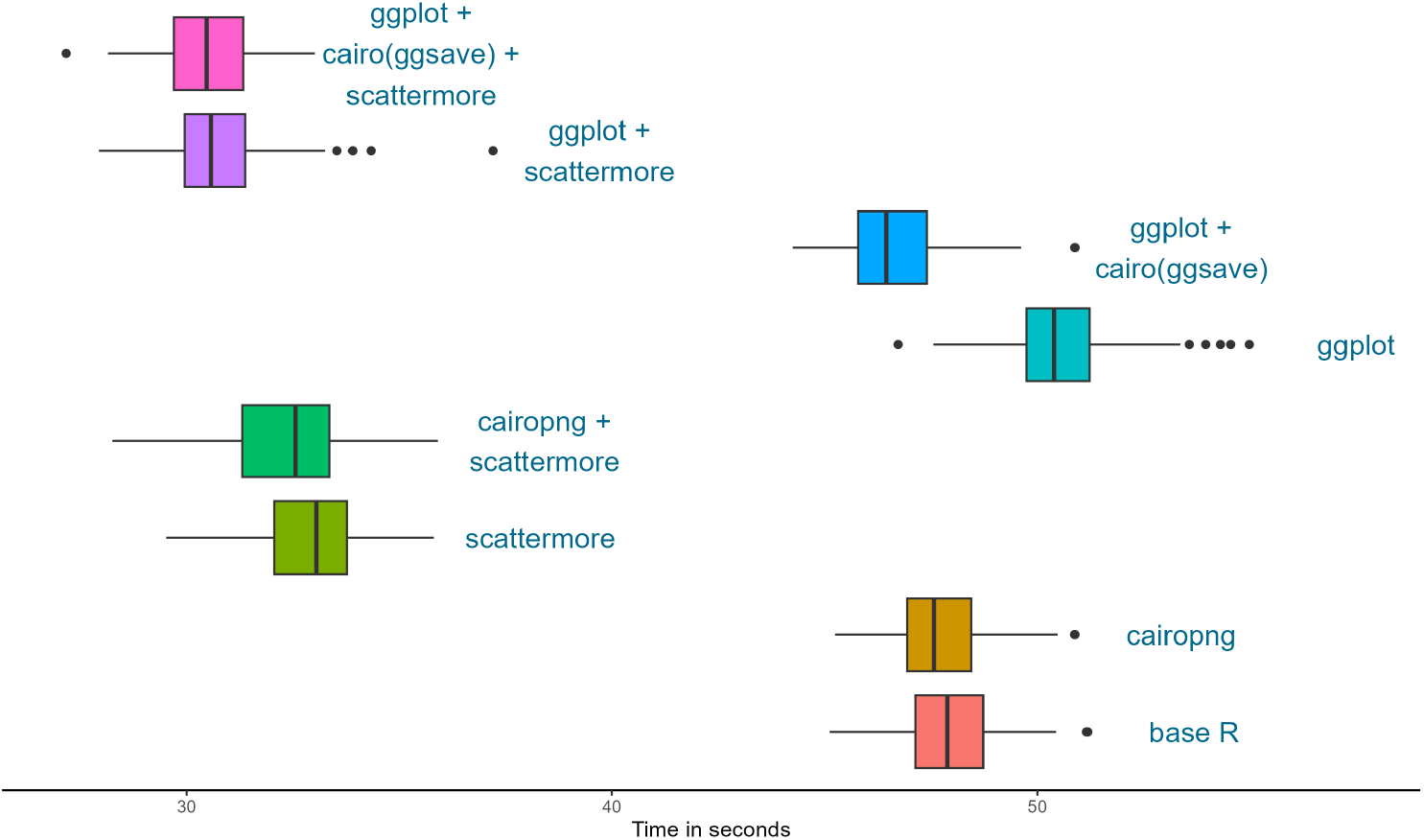
Time comparison among different `fastman()` plot-generating options.

As we can see, using **scattermore** reduces the plotting time substantially. The combination of **scattermore** and **ggplot** produces the fastest plots. The package converts data points to a raster layout before plotting, saving considerable time during plotting. However, the rasterisation step needs pre-specification of plotting areas for optimal display. Hence it can look grainy or blurry if the specification is not properly in sync with the plot size parameters while saving the plot. The parameters used in our function represent an optimal middle ground between visual acuity and speed. However, if the user wants the best visual display it is better not to use **scattermore**; only use it if speed is of the highest importance.

We now compare the `fastman()` base R plot with the existing packages (**qqman** (Turner 2018) and **fastman (Daniel)** (Hwang 2016)) in terms of time taken for plot generation. There are many other packages for Manhattan plot visualisation of GWAS results, such as **geni.plots** (Staley 2024), which provide various additional flexibilities but do not specifically focus on faster plotting; therefore for simplicity, we did not assess their performance in our comparisons. For this comparison, we used the same data set from Figure 4 of 8,776,402 rows. We ran a hundred iterations on this data set. The median time taken by the packages across the iterations has been reported in Figure 12.

**Figure 12:**
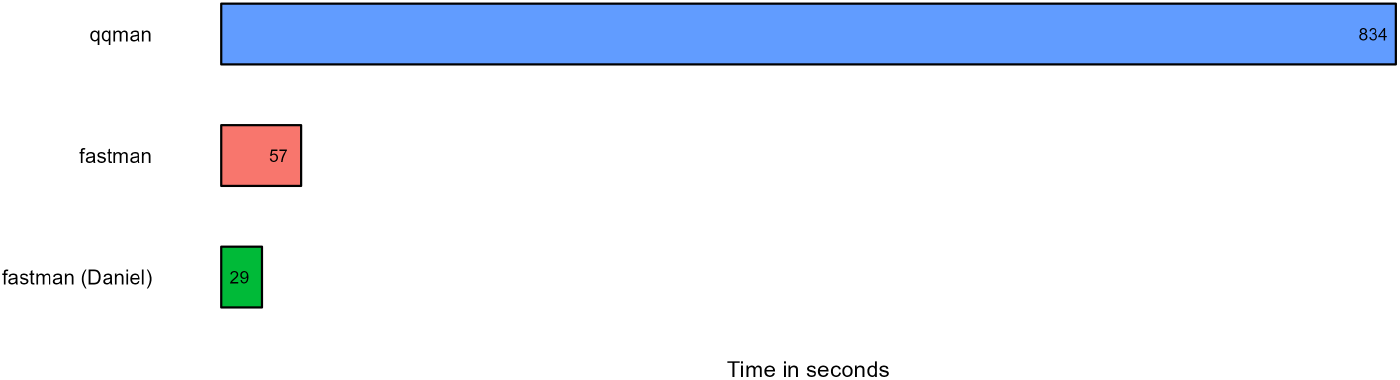
Time comparison of **fastman** with other packages.

We observe that **fastman** reduces plot generation time drastically compared to **qqman. fastman** (Daniel) is marginally faster than **fastman** as it restricts the number of SNPs with p-values between 0.01 and 0.1 to 20000, and hence has a much smaller data set to generate a plot from. We expect the package also to show similar behaviour in other data sets.

Looking more closely at the different stages of code execution, in Figure 13 (A) we see that **fastman** takes marginally more time than **qqman** while preparing the data for plotting, during the ‘initial calculations’ and ‘plotting positions’ stages, but it is orders of magnitude faster during the actual plot generation step (having reduced it to a smaller data), which is the most time-consuming step in **qqman;** hence **fastman** is much faster overall.

**Figure 13:**
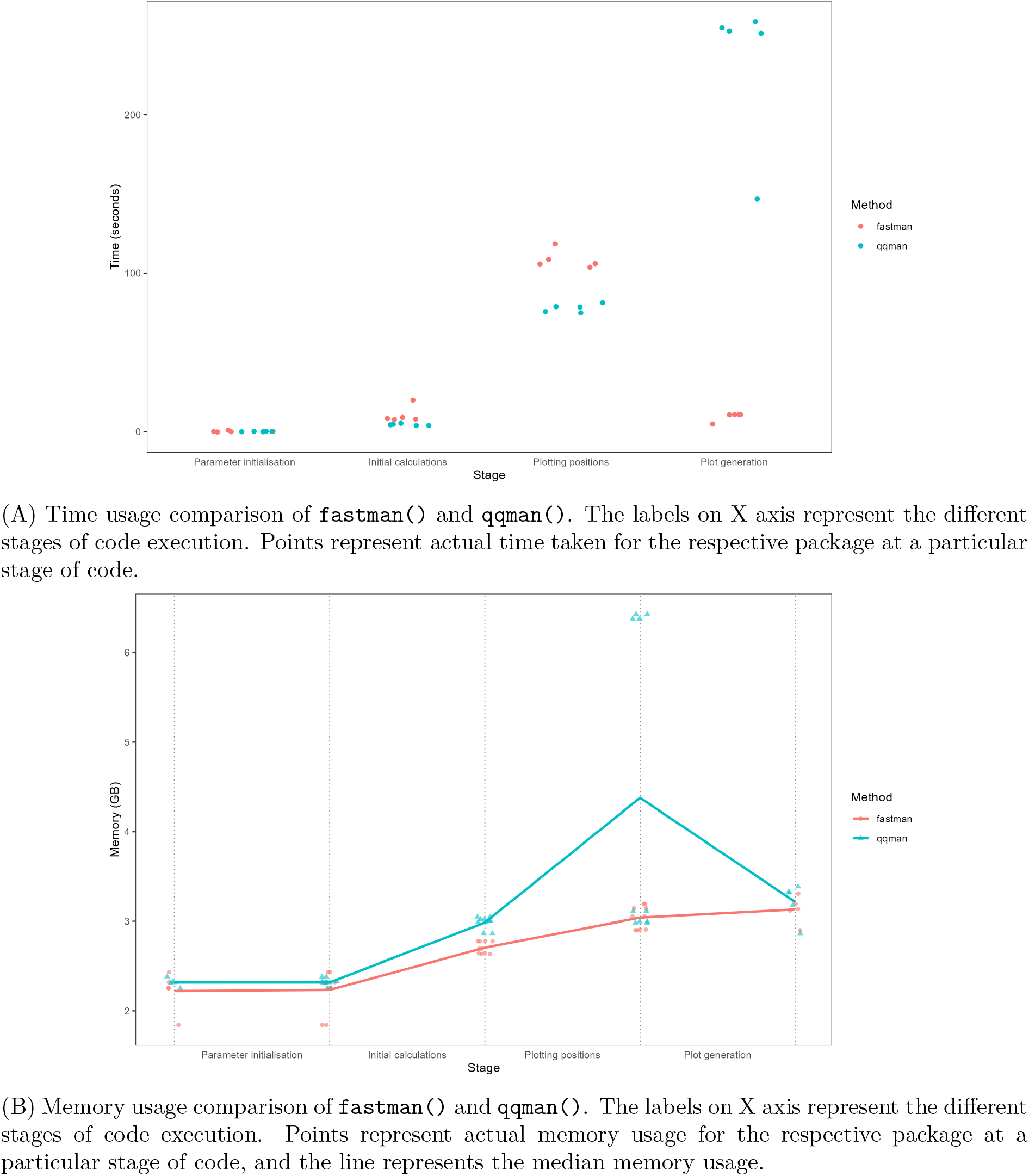
Time and memory usage comparison of `fastman()` and `qqman()`.

Using the same data set, we compared the memory usage of **fastman** and **qqman** across different phases of execution, from parameter initiation, interim calculations and adjustments to the final plot generation. As demonstrated in Figure 13 (B), **fastman** utilises lower memory space than **qqman** despite the additional calculations used for speeding up. Each dotted line represents the start or end of a specific stage of the code execution. The memory usage gradually increases as the code reads the data and prepares the data for plot generation.

As described previously, **fastman** reduces the plotting time by reducing the final number of data points to be plotted. To demonstrate this, we used the dataset from Figure 4 of 878k rows. Going from 1% to 100% of the number of rows in the data, we selected random subsets of rows and created Manhattan plots using **fastman**. For every percentile, we took 10 random subsets, performed plotting, and recorded the plotting times and final number of data points plotted. Figure 14 shows how the plotting time of **fastman** varies with the number of data points. We can see that plotting time increases linearly as the number of rows increases.

**Figure 14:**
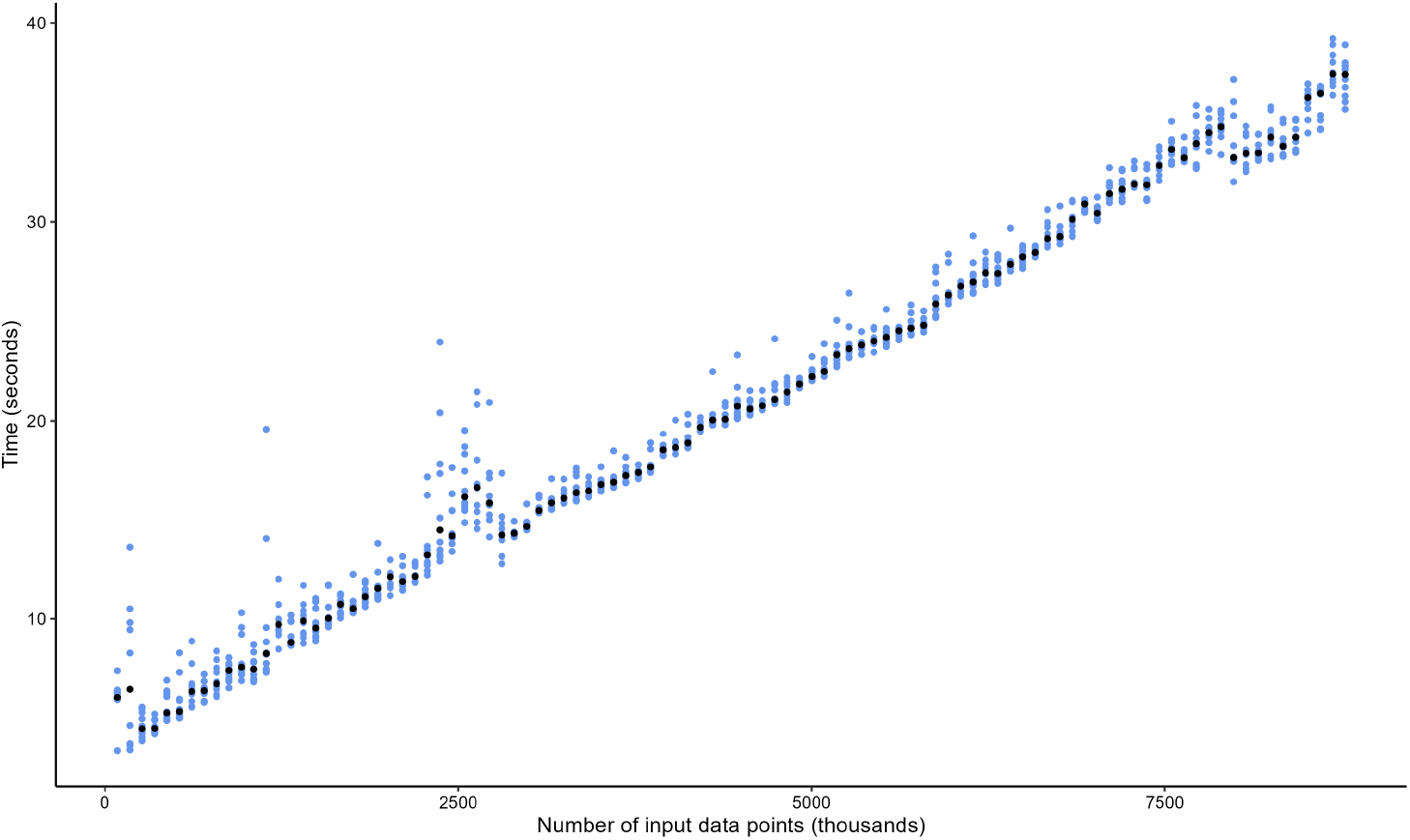
Time taken by `fastman()` as the number of data points in the input data frame increases. For a particular input data size, the black points represent the median time taken, while the blue points represents the actual time taken for plot generation. The median points show an approximately linear relationship with input data size.

The next Figure 15 shows the final number of data points plotted by **fastman** against the number of data points initially. We can observe a logarithmic relation between the initial and final number of data points plotted by fastman. We have fitted a *y* = −189.22 + 58.76 log(*x*) regression line (marked with red) in the plot to demonstrate the said relationship. Given the logarithmic nature of the number of final data rows plotted, it becomes clear that the speedup advantage is the highest for larger datasets.

**Figure 15:**
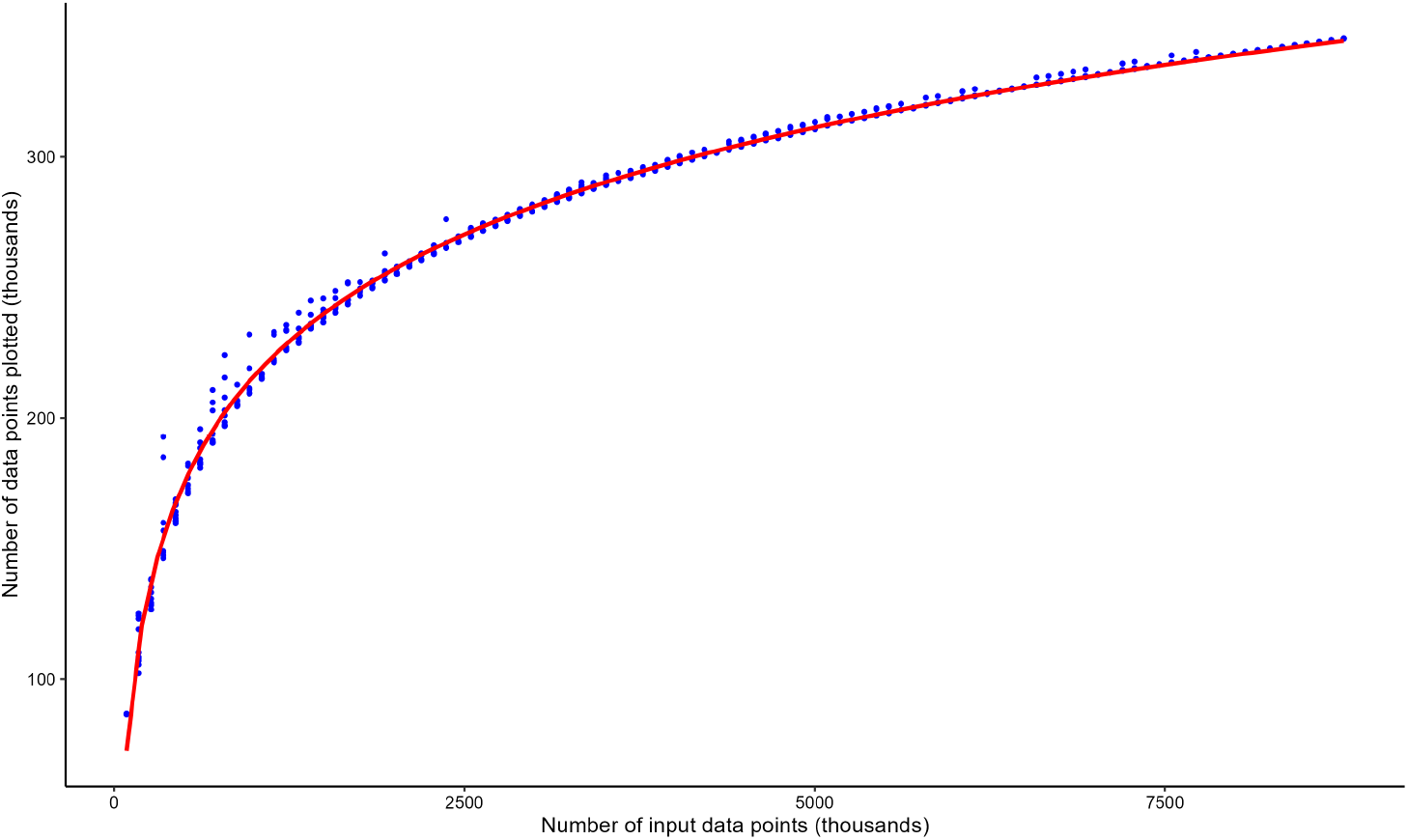
Number of final data points plotted by `fastman()` as number of data points in the input data frame increases. The fitted *y* = −191.54 + 59.03 log(*x*) regression line marked in red demonstrates the logarithmic relation between the initial and final number of data points plotted.

### 4.3. Performance Comparison: fastqq

To compare fastqq() with existing QQ plot packages and functions, we used the same data set of 8,776,402 rows from Figure 4. We ran ten iterations for all the packages/functions, and the median times in seconds are plotted in Figure 16.

**Figure 16:**
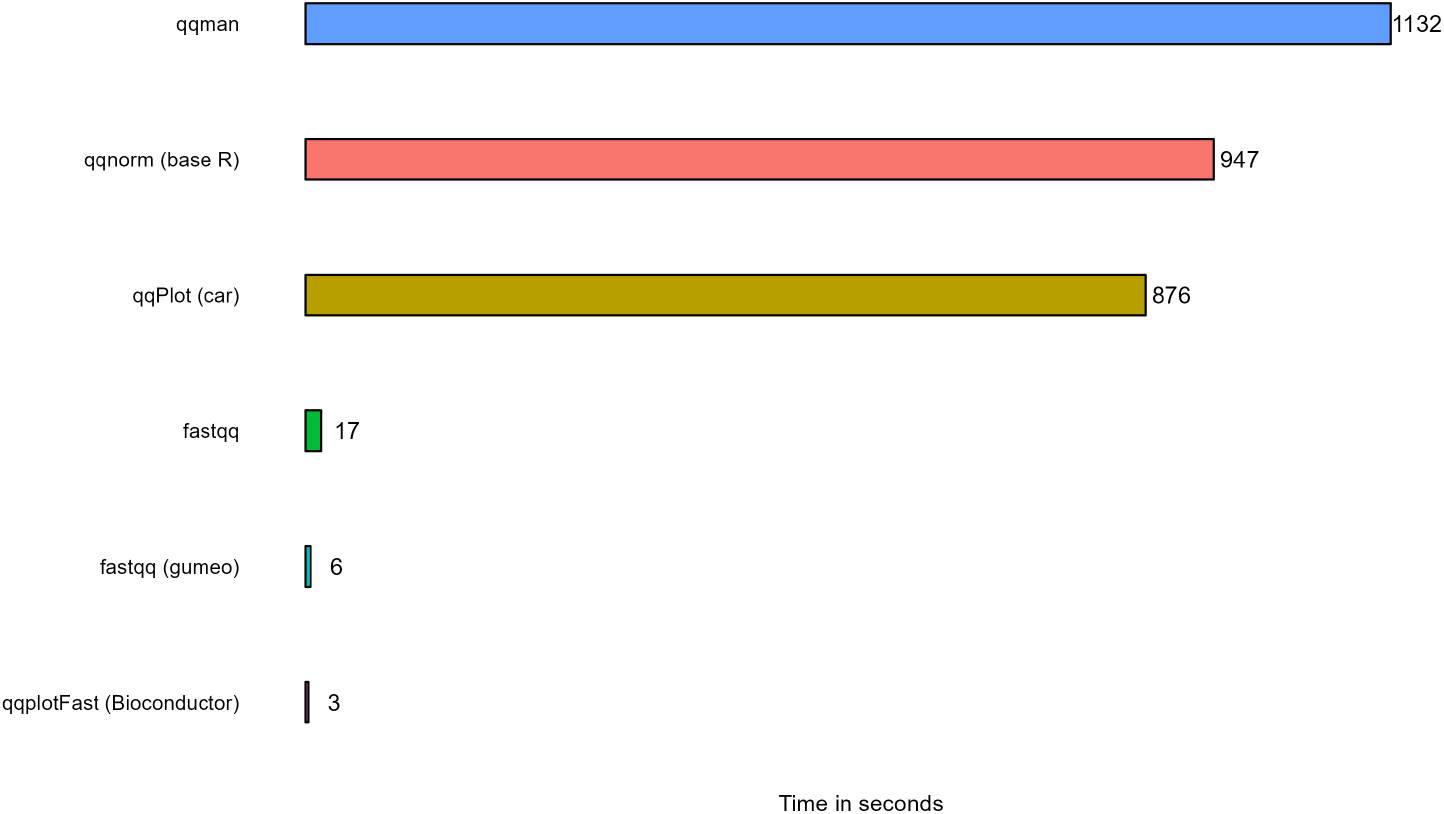
Time comparison of `fastqq()` with other packages.

Our function fastqq() reduces plotting time drastically compared to **qqman**, the qqPlot() function in **car** package, and the base R qqnorm() function. Only qqplotFast() function from **raMWAS** (Shabalin et al. 2018) package in **Bioconductor** and **fastqq (gumeo)** (Einarsson and Einarsson 2022) are observed to be marginally faster than fastqq().

## 5. Summary and Discussion

**fastman** is an R package that offers the user a fast and efficient option for visualising GWAS results using Q-Q and Manhattan plots. While this can also be accomplished using various other software packages, **fastman** manages to reduce plotting time without any data loss and hence does not compromise on the exactness of the visualisation. It also provides the user with many input flexibility and output customisation options while drastically reducing the plot generation time. The ability to work with non-model organisms is particularly relevant for researchers working in many other disciplines. The future steps in the **fastman** package will include adding additional features, such as a function for Miami plots (which shows two Manhattan plots for two different GWAS results, plotted along the same X axis representing genomic locations, with one set of values being plotted along the positive y-axis and the other along the negative side), implementing the package in Python, and integrating it with popular R genomic packages.

## Acknowledgments

SSP and KA wrote the R scripts, designed the package and maintained the GitHub repository. SSP, SRR and KA did the testing for different data types. SSP performed the performance comparisons, prepared the figures and drafted the manuscript. SRR and KA revised, edited and made relevant additions to the manuscript. All authors proofread and approved the final manuscript. We are grateful to Stephen Turner for creating **qqman** and encouraging us during the development of this package. SRR thanks Jeffrey D. Lozier and Heather M. Hines for their valuable input and suggestions. SSP is thankful to Sagnik Palmal for being one of the earliest users of our package. We express our sincerest gratitude to all **fastman** users who have performed additional tests of the package, reported bugs, and made useful suggestions that we have incorporated into the final package.

## Appendix

### A. Other Statistics for Visualisation

Apart from the negative logarithm of P-values from GWAS, Manhattan plot can be used for visualising several other genome-wide population genetic parameters.

#### A.1. Fixation Index

Fixation index or *F*_*ST*_ (Holsinger and Weir 2009) is a measure of genetic differentiation between populations. It is based on the variance in allele frequencies between populations relative to the total variance across populations. *F*_*ST*_ ranges from 0 to 1:

- *F*_*ST*_ = 0 indicates no genetic differentiation (populations are genetically identical)
- *F*_*ST*_ = 1 indicates complete genetic differentiation (the populations do not share any alleles)

This statistic is often used to assess the degree of population structure or isolation, and higher *F*_*ST*_ values suggest more genetic differentiation between populations. It is widely used in studies of population genetics to evaluate the level of genetic variation within and between subpopulations.

Mathematically, *F*_*ST*_ can be calculated using the following formula:

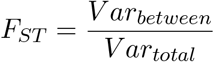

where *V ar*_*between*_ is the variance in allele frequencies between populations, and *V ar*_*total*_ is the total genetic variance across the entire population.

When visualised using Manhattan plots, *F*_*ST*_ values are typically plotted across genomic regions, and loci that exhibit significant differentiation between populations are highlighted for further inspection.

#### A.2. Nucleotide Diversity

*π* statistics (Nei and Li 1979) is a measure of genetic diversity within a population. It represents the average number of nucleotide differences per site between two randomly chosen sequences from the same population. This statistic is typically used to quantify the extent of genetic variation present in a population.

*π* is calculated as the average pairwise differences between all pairs of sequences (or individuals) in a population. Higher *π* values indicate more genetic diversity.

Mathematically, *π* can be calculated as:

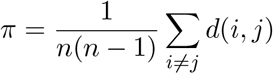

where *n* is the number of sequences or individuals, and *d*_*ij*_ is the number of differences between sequences *i* and *j*.

In a Manhattan plot, *π* can be visualised across the genome, with each point representing a genomic region and showing the variation in genetic diversity. This can be used to identify regions with high or low genetic diversity.

#### A.3. D-statistics

D-statistics (Green et al. 2010), often referred to as the ABBA-BABA test, is a statistic used to test for introgression or gene flow between populations. This test is particularly useful when comparing the genomic data of three populations or species and looking for evidence of genetic admixture, where one population has contributed genes to another.

The D-statistic compares two specific types of allele-sharing patterns:

- **ABBA pattern**: One population shares alleles with another, while a third population (often an outgroup) shares the other allele
- **BABA pattern**: The reverse occurs, where the allele-sharing pattern is inverted between the populations

The D-statistic is calculated as:

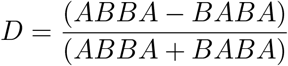

where ABBA refers to sites where the alleles are shared between populations A and B, but not the outgroup, and BABA refers to sites where the alleles are shared between populations B and the outgroup, but not population A.

A significant positive or negative D-statistic provides evidence for gene flow or introgression between populations, with values significantly different from zero suggesting the presence of admixture.

When plotted on a Manhattan plot, D-statistics can highlight regions where significant introgression or gene flow events have occurred. This can be observed by looking at substantial deviations from expected allele frequency patterns.

### B. Validation of Chosen Threshold for Uniform Distribution

We are deriving the distribution of the statistics *R* and *L* in case of uniform distribution. We are going to show that, for uniform distribution, the probability of *R* and *L* exceeding the threshold 0.1 is equal to 0, thereby validating our choice of the threshold. While a closed form theoretical distribution of these statistics are possible to derive in case of uniform, this is not possible for other distributions like normal or exponential.

#### B.1. Distribution of R

We need to find the distribution of the ratio

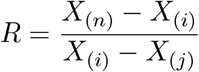

where *X*_(*j*)_, *X*_(*i*)_, and *X*_(*n*)_ are the order statistics of a uniform(0,1) sample of size *n*, with *j < i < n*.

##### Step 1: Joint distribution of order statistics

The joint density of *X*_(*j*)_, *X*_(*i*)_, and *X*_(*n*)_ for a uniform(0,1) sample is given by:

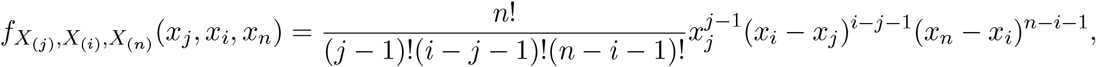

for 0 ≤ *x*_*j*_ ≤ *x*_*i*_ ≤ *x*_*n*_ ≤ 1.

###### Derivation

To compute this, we need to count how many of the original *n* observations fall in the three regions:

- *j* − 1 values are less than *x*_*j*_
- *i* − *j* − 1 values are between *x*_*j*_ and *x*_*i*_
- *n* − *i* − 1 values are between *x*_*i*_ and *x*_*n*_

The probability of this happening is obtained by considering the probability of selecting these values and arranging them in this order. So, we can write:

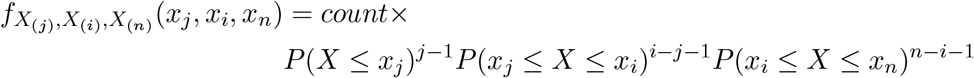

where *X* ~ Unif (0, 1), and *count* is the number of ways these points can be arranged. This is given by:

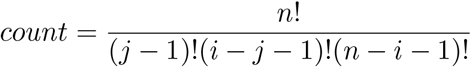

We also know that, if *X* ~ Unif (0, 1), then:

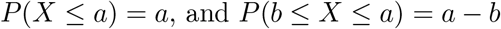

Combining these two, we get:

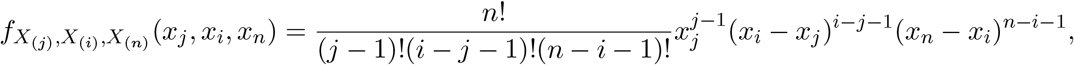

##### Step 2: Variable transformation

Let us define the transformed variables:

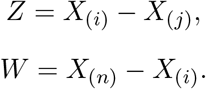

Thus, we are interested in the following ratio:

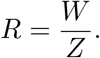

Rewriting the order statistics in terms of *Z* and *W* :

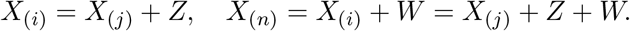

The Jacobian determinant for this transformation is:

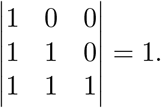

So the joint density remains:

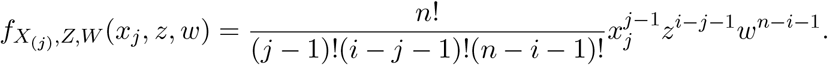

##### Step 3: Distribution of *R*

To find *f*_*R*_(*r*), we marginalise over *X*_(*j*)_. Since the integration limits are 0 ≤ x_j_ ≤ 1 − (*z* + *w*), we integrate out *x*_*j*_:

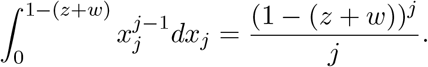

So the joint density of *Z* and *W* is:

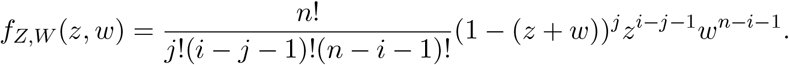

Now we change variables to *Z* and *R* = *W/Z*, so *W* = *RZ* and *dW* = *ZdR*. The density transformation gives:

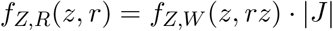

where the Jacobian determinant is:

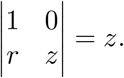

Substituting *w* = *rz*, we get:

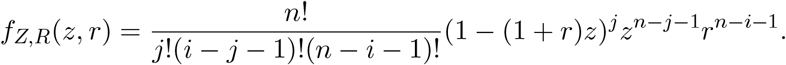

Now we integrate over *z* from 0 to 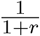 to get *f*_*R*_(*r*):

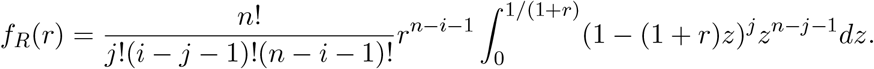

Let us consider this transformation: *t* = (1 + *r*)*z*, then the density becomes:

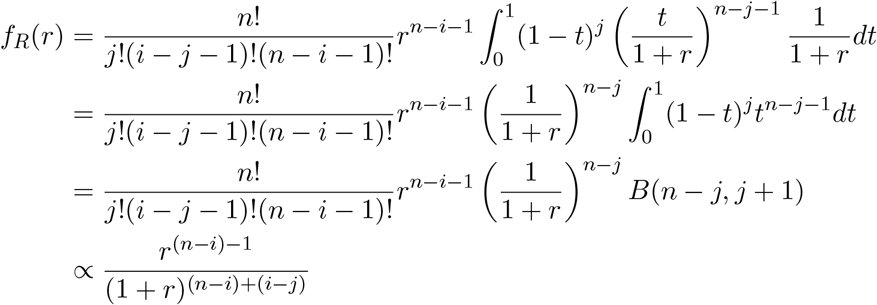

Comparing this with the standard Beta Prime distribution, which has the form:

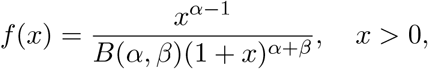

we identify the parameters as

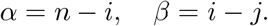

Thus, *R* follows a Beta Prime distribution:

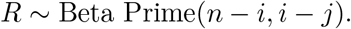

In our case, *R* has been defined as

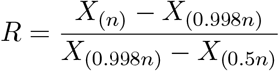

Hence, we can say that

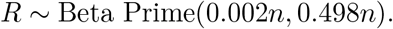

Or equivalently

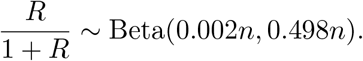

For our choice of threshold 0.1, we can show that R is always less than this threshold by showing:

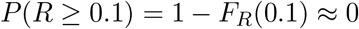

The CDF of Beta Prime distribution is the incomplete Beta distribution, which does not have a closed form expression. So we have evaluated this numerically for several values of *n* between 1,000 and 1,000,000. The value, reported by R (software), implemented in `1 − pbeta(0.1 / (1 + 0.1), 0.002*n, 0.498*n)` was returned to be exactly 0 up to the numerical tolerance of R (software).

#### B.2. Distribution of L

We need to find the distribution of the ratio

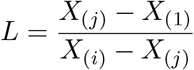

where *X*_(1)_, *X*_(*j*)_, and *X*_(*i*)_ are the order statistics of a uniform(0,1) sample of size *n*, with *i > j >* 1.

##### Step 1: Joint distribution of order statistics

The joint density of *X*_(1)_, *X*_(*j*)_, and *X*_(*i*)_ for a uniform(0,1) sample is given by:

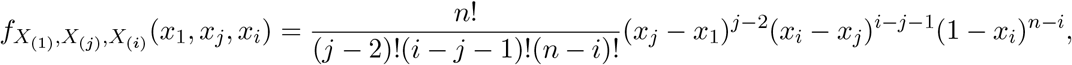

for 0 ≤ *x*_*j*_ ≤ *x*_*i*_ ≤ *x*_*n*_ ≤ 1.

This can be derived in a very similar manner like *R*. We need to count how many of the original *n* observations fall in the three regions:

- *j* − 2 values are between than *x*_1_ and *x*_*j*_
- *i* − *j* − 1 values are between *x*_*j*_ and *x*_*i*_
- *n* − *i* values are greater than *x*_*i*_

##### Step 2: Variable transformation

Let us define the transformed variables:

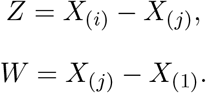

Thus, we are interested in the following ratio:

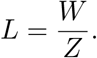

Rewriting the order statistics in terms of *Z* and *W* :

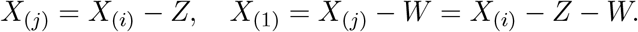

The Jacobian determinant for this transformation is:

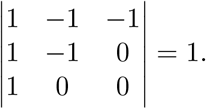

So the joint density remains:

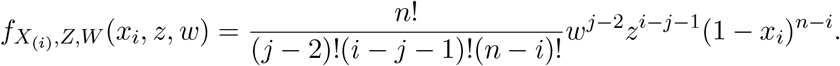

##### Step 3: Distribution of *L*

To find *f*_*L*_(*l*), we marginalise over *X*_(*i*)_. Since the integration limits are (*z* + *w*) ≤ x_i_ ≤ 1, we integrate out *x*_*i*_:

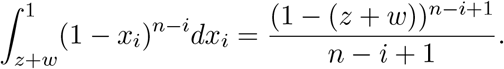

So the joint density of *Z* and *W* is:

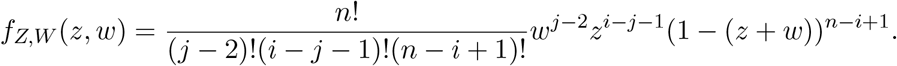

Now we change variables to *Z* and *L* = *W/Z*, so *W* = *LZ* and *dW* = *ZdL*. The density transformation gives:

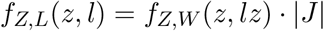

where the Jacobian determinant is:

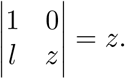

Substituting *w* = *lz*, we get:

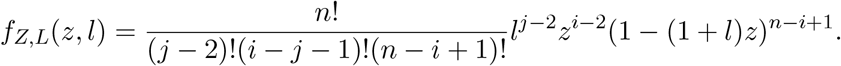

Now we integrate over *z* from 0 to 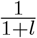 to get *f*_*L*_(*l*):

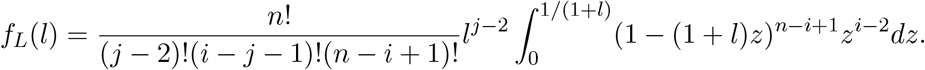

Let us consider this transformation: *t* = (1 + *l*)*z*, then the density becomes:

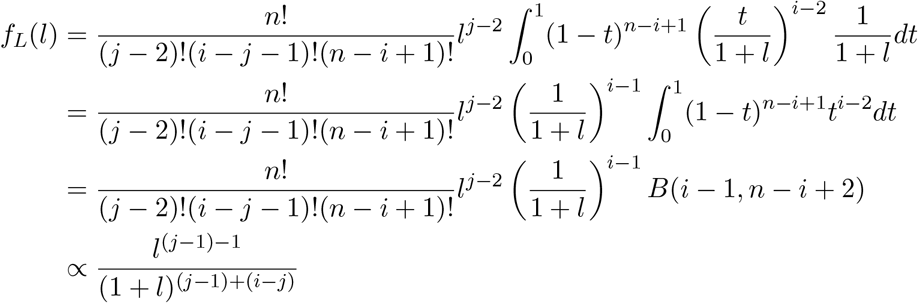

Comparing this with the standard Beta Prime distribution, which has the form:

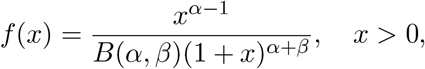

we identify the parameters as

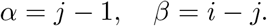

Thus, *L* follows a Beta Prime distribution:

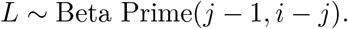

In our case, *L* has been defined as

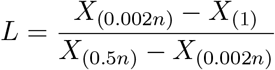

Hence, we can say that

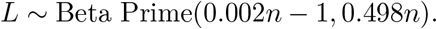

Or equivalently

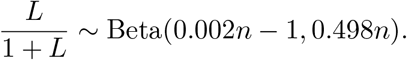

For our choice of threshold 0.1, we can show that L is always less than this threshold by showing:

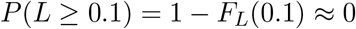

Just like the case of statistic *R*, we have evaluated *P* (*L* ≥ 0.1) numerically for several values of *n* between 1,000 and 1,000,000, as the CDF of Beta Prime distribution does not have a closed form expression. The value, reported by R (software), implemented in `1 − pbeta(0.1 / (1 + 0.1), 0.002*n-1, 0.498*n)` was returned to be exactly 0 up to the numerical tolerance of R (software).

